# Dual airway and alveolar contributions to adult lung homeostasis and carcinogenesis

**DOI:** 10.1101/531780

**Authors:** Magda Spella, Ioannis Lilis, Mario A. Pepe, Yuanyuan Chen, Maria Armaka, Anne-Sophie Lamort, Dimitra E. Zazara, Fani Roumelioti, Malamati Vreka, Nikolaos I. Kanellakis, Darcy E. Wagner, Anastasios D. Giannou, Vasileios Armenis, Kristina A.M. Arendt, Laura V. Klotz, Dimitrios Toumpanakis, Vassiliki Karavana, Spyros G. Zakynthinos, Ioanna Giopanou, Antonia Marazioti, Vassilis Aidinis, Rocio Sotillo, Georgios T. Stathopoulos

**Author notes:** Equal senior co-authors (Rocio Sotillo, Georgios T. Stathopoulos). Corresponding authors: Magda Spella, PhD and Georgios T. Stathopoulos, MD PhD; Laboratory for Molecular Respiratory Carcinogenesis, Department of Physiology, Faculty of Medicine, University of Patras; Basic Biomedical Sciences Building, 2nd Floor, Room B40, 1 Asklepiou Str., University Campus, 26504 Rio, Greece; and; and Lung Carcinogenesis Group, Comprehensive Pneumology Center (CPC) and Institute for Lung Biology and Disease (iLBD), Max-Lebsche-Platz 31, 81377 Munich, Germany.

## Abstract

Lung adenocarcinoma (LUAD) and chronic lung diseases caused by smoking and environmental noxious agents are the deadliest diseases worldwide, sharing a partially charted pathobiology of dysfunctional alveolar repair. Here we sought to identify the respiratory epithelial dynamics and molecular signatures participating in adult lung maintenance and chemical carcinogenesis. We employed novel mouse models of respiratory epithelial marking and ablation, a battery of pulmonary toxins and carcinogens, experimental protocols of carcinogen-induced LUAD, tobacco carcinogen-induced LUAD cell lines, and human transcriptomic data and identified a prominent involvement of airway molecular programs in alveolar maintenance and carcinogen-induced LUAD. The airway-specific transcriptomic signature was redistributed to the alveoli after toxic and carcinogenic insults and resulted in marked contributions of airway-labeled cells to injury-recovered alveoli and LUAD. Airway cells maintained *Kras* mutations and therefore possibly contributed to lung cancer initiation, while LUAD were spatially linked to neighboring airways. Transcriptomic profiling of carcinogen-induced murine and human LUAD revealed enrichment in airway signatures, while ablation of airway cells distorted alveolar structure and function and protected mice from LUAD development. Collectively, these results indicate that airway cells and/or transcriptomic signatures are essential for alveolar maintenance and LUAD development.

## INTRODUCTION

Chronic lung diseases present tremendous health burdens attributed to dysfunctional alveolar repair [1]. Lung adenocarcinoma (LUAD), the leading cancer killer worldwide, is mainly caused by chemical carcinogens of tobacco smoke that induce mutations of the Kirsten rat sarcoma viral oncogene homologue (*KRAS*) in distal pulmonary cells [2–4]. The identification of the cellular and transcriptomic events that underlie lung regeneration and carcinogenesis is extremely important, since epithelial developmental pathways are intimately related with oncogenic signaling to jointly regulating stemness and drug resistance [5, 6]. To this end, lineage-specific genes encoding epithelial proteins that support the physiological functions of the lungs were recently shown to suffer non-coding insertions and deletions in LUAD, lending further support to the longstanding notion that epithelial cells that express lung-restricted proteins are the cellular sources of LUAD [7].

Previous pulmonary lineage tracing studies that utilized noxious insults and expression of oncogenic *KRAS* in the respiratory epithelium incriminated different cells as progenitors of new alveoli and/or LUAD in adult mice: airway epithelial cells (AEC) including basal and Clara or club cells, alveolar type II cells (ATII), and/or bronchoalveolar stem cells with dual ATII/Clara properties [8–14]. However, mouse lineage tracing models feature incomplete/promiscuous lung cell labeling [8–14]. In addition, the mutational landscape of human LUAD is closely mirrored by tobacco carcinogen-induced, but not by oncogenic *KRAS*-driven mouse lung tumors [15].

We combined accurate genetic marking of airway and alveolar epithelial cells with different insults to the adult lung to integrally assess the signatures that contribute to alveolar maintenance, repair, and carcinogenesis. This approach was adopted to identify the molecular signature of regenerating and malignant cells, which was recently shown to be linked with mutation processes [7]. To achieve our goals, we adapted multi-hit chemical carcinogen exposure protocols to the murine *C57BL/6* strain that is resistant to chemical tumor induction [16, 17], and corroborated the findings with the *FVB* strain that is susceptible to single-hit carcinogenesis [15, 18]. We show that in the adult mouse, aging-, pneumotoxin- and carcinogen-insults result in a marked activation of airway transcriptomic programs across the alveolar parenchyma, contributing to alveolar repair and carcinogenesis. We determined that, early after chemical carcinogen exposure, airway cells preferentially suffer and sustain *Kras* oncogenic mutations compared with alveolar cells; that LUAD are spatially associated with neighboring airways; and that ablation of airway cells hinders alveolar maintenance and carcinogenesis in mice. The data indicate remarkable plasticity of adult lung epithelial cells and suggest a central role for airway cells and transcriptional signature sin lung regeneration and cancer.

## RESULTS

### Accurate genetic marking of airway and alveolar cells

For genetic marking, we crossed a *Cre*-reporter strain that features *Cre*-mediated permanent switch of somatic cells from membranous tdTomato (*mT*) to GFP (*mG*) fluorescence (*mT/mG*) to six different *Cre*-driver strains, including a novel airway-specific *Cre*-driver (*Scgb1a1.Cre*, *Sftpc.Cre*, *Lyz2.Cre*, *Sox2.Cre*, *Vav.Cre*, and *Nes.Cre*; all *C57BL/6* background; [14, 19–24]). In double heterozygote offspring at six postnatal weeks (i.e., after lung development is complete; [8, 14]), *mG+* labeling was completely and exclusively confined to the airway epithelium of *mT/mG;Scgb1a1.Cre* mice, promiscuous to airway and alveolar epithelia of *mT/mG;Sftpc.Cre* mice, partial and exclusive to the alveolar epithelium of *mT/mG;Lyz2.Cre* mice, and not informative in the remaining intercrosses (Figure 1A and Figure S1). Co-localization of *mG+* labeling with lineage marker proteins revealed complete *mG+* labeling of all AEC but not of alveolar cells in *mT/mG;Scgb1a1.Cre* mice, of most AEC and all ATII cells in *mT/mG;Sftpc.Cre* mice, and of some ATII cells and all alveolar macrophages in *mT/mG;Lyz2.Cre* mice (Figure 1A and Figures S1B and S1C). Lung flow cytometry of six-week-old *mT/mG;Scgb1a1.Cre*, *mT/mG;Sftpc.Cre*, and *mT/mG;Lyz2.Cre* mice estimated the proportions of *mG*+ marked cells in concordance to microscopy (Figure 1A and Figures S1A and S1D). Thus *mT/mG;Scgb1a1.Cre* and *mT/mG;Lyz2.Cre* mice display 100% distinct labeling of AEC versus ATII cells and AMΦ at conclusion of development.

**Figure 1.**
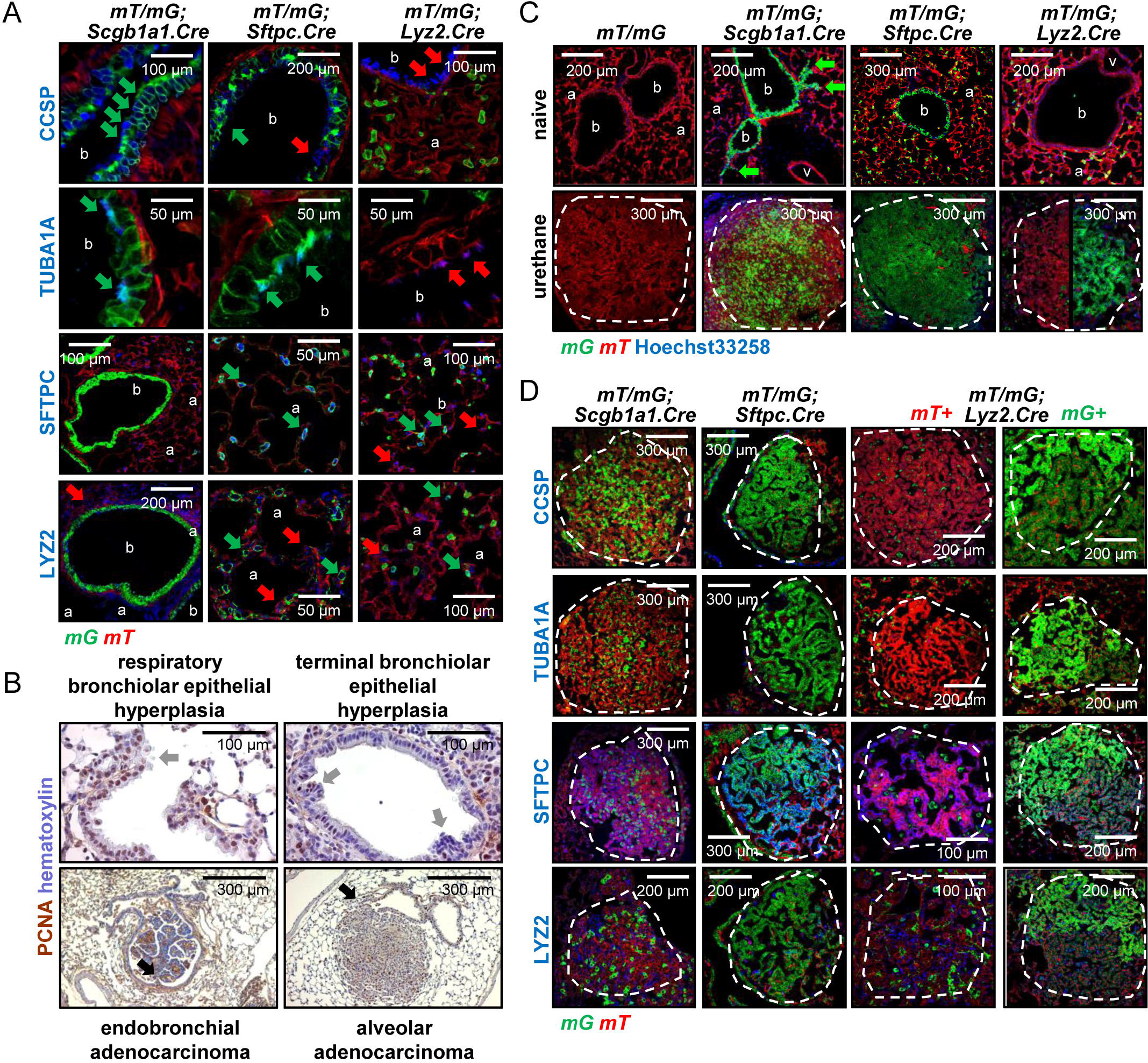
Multiple epithelial signatures in urethane-induced lung adenocarcinoma (LUAD). **A,** Lineage marker-stained lung sections of 6-week-old marked mice (*n* = 5/group). Note *mG*+CCSP+ Clara and *mG*+TUBA1A+ ciliated cells in bronchi (b) of *mT/mG;Scgb1a1.Cre* mice, *mG*+SFTPC+LYZ2± alveolar type II cells and *mG*+SFTPC-LYZ2+ alveolar macrophages in the alveoli (a) of *mT/mG;Lyz2.Cre* mice, and various *mG*+ cells in bronchi and alveoli of *mT/mG;Sftpc.Cre* mice. Arrows: Lineage marker protein-expressing *mG*+ (green) and *mT*+ (red) cells. **B,** Proliferating cell nuclear antigen (PCNA)-stained lung sections of urethane-treated *C57BL/6* mice at six months into treatment. Arrows: airway hyperplasias (grey) and LUAD (black). **C,** Lungs of genetically marked mice before (top; *n* = 5/group) and six months after (bottom; *n* = 30, 22, 18, and 20/group, respectively) urethane commencement. Note *mG*+ cells in terminal and respiratory bronchioles (arrows) and LUAD (dashed outlines) of *mT/mG;Scgb1a1.Cre* mice, ubiquitous *mG*+ airway, alveolar, and LUAD cells of *mT/mG;Sftpc.Cre* mice, and partial alveolar and LUAD marking of *mT/mG;Lyz2.Cre* mice. v, vessel. **D,** Lineage marker protein-stained LUAD (dashed outlines) from genetically marked mice from C (*n* = 10/group). Note *mG*+CCSP-SFTPC+LYZ2± LUAD cells of *mTmG;Scgb1a1.Cre* mice. Five non-overlapping fields/sample were examined. *mG*, membranous green fluorescent protein fluorophore; *mT*, membranous tomato fluorophore; CCSP, Clara cell secretory protein; TUBA1A, acetylated α-tubulin; SFTPC, surfactant protein C; LYZ2, lysozyme 2.

### Airway and alveolar signatures in chemical-induced lung adenocarcinoma

We reproducibly induced LUAD in the above genetically-marked mice using repetitive exposures to the tobacco-contained carcinogens urethane (ethyl carbamate, EC; stand-alone mutagen and tumor promoter) or 3-methylcholanthrene followed by butylated hydroxytoluene (MCA/BHT; a two-hit mutagen/tumor promoter regimen; Figure S2A). In both models, preneoplastic (airway epithelial hyperplasia, atypical alveolar hyperplasia) and neoplastic (adenoma and LUAD) lesions [25] were both airway- and alveolar-located and hence inconclusive on tumor origins (Figure 1B). Interestingly, both airway- and alveolar-located hyperplasias and tumors of *mT/mG;Scgb1a1.Cre* mice were partially *mG*+, of *mT/mG;Sftpc.Cre* ubiquitously and non-informatively *mG*+, and of *mT/mG;Lyz2.Cre* mice either *mG*+ or *mG*- (Figure 1C and Figure S2B-S2E). Immunostaining revealed that genetically-marked *Scgb1a1+*, *Sftpc+*, and *Lyz2+* tumor cells were CCSP-TUBA1A-SFTPC+LYZ2± (Figure 1D). We further tested these observations using a one-hit model in which a single exposure of susceptible *FVB* mice to urethane causes LUAD featuring the full-blown mutation spectrum of the human disease [15, 18]. For this, we backcrossed *mT/mG*, *Scgb1a1.Cre*, *Sftpc.Cre* and *Lyz2.Cre* strains >F12 to the *FVB* background, set up all the relevant intercrosses, and subjected double heterozygote offspring to a single urethane exposure. Examination of neoplastic lesions at six months after carcinogen again revealed mixed contributions of both airway and alveolar cells/signatures to LUAD (Figure S3A-S3C). Baseline genetic marking at six postnatal weeks was similar to that observed in the respective genotypes in the *C57BL/6* strain (*mG*+ marking in Figure S3D), as was the expression pattern of specific lineage markers (Figure S3D), corroborating ubiquitous marking of all CCSP+ AEC in *mT/mG;Scgb1a1.Cre* mice, of both CCSP+ AEC and SFTPC+ ATII cells in *mT/mG;Sftpc.Cre* mice, and of some SFTPC+ ATII cells in *mT/mG;Lyz2.Cre* mice. Collectively, these data support that chemical-induced LUAD carry both airway and alveolar signatures and suggest that tumor cells can originate from *Scgb1a1+* airway cells that up-regulate SFTPC with or without LYZ2 during carcinogenesis, from *Scgb1a1+Sftpc+* airway cells that feature CCSP loss with/without LYZ2 gain, or from *Sftpc+* alveolar cells that transiently express CCSP with/without LYZ2.

### Airway cells sustain *Kras* mutations

We next determined the lung lineage in which LUAD driver mutations are inflicted. Since urethane-induced LUAD of mice exclusively harbor *Kras*^Q61R^ mutations [15], we sought these at early time-points after urethane. For this, *FVB mT/mG;Scgb1a1.Cre* and *mT/mG;Lyz2.Cre* mice received urethane, lungs were harvested one and two weeks later, and digital droplet PCR was performed with probes targeting *mT* and *Kras*^Q61R^ sequences. Interestingly, *mG*+;*Kras*^Q61R^ cells in the lungs of *mT/mG;Scgb1a1.Cre* mice (i.e. *Kras*^Q61R^-mutant AEC) survived and increased in number, while *mG*+;*Kras*^Q61R^ cells in the lungs of *mT/mG;Lyz2.Cre* mice (i.e. *Kras*^Q61R^-mutant ATII cells) did not persist over time (Figure 2A). Supporting the importance of AEC in LUAD development, three-dimensional reconstruction of chemical carcinogen-inflicted lungs of *FVB* mice using high-resolution micro-computed tomography (μCT) revealed that most lung tumors were spatially linked with neighboring airways, either sprouting from or even contained in bronchi (Figure 2B). These results support a significant role of airway cells in chemical-induced LUAD of mice.

**Figure 2.**
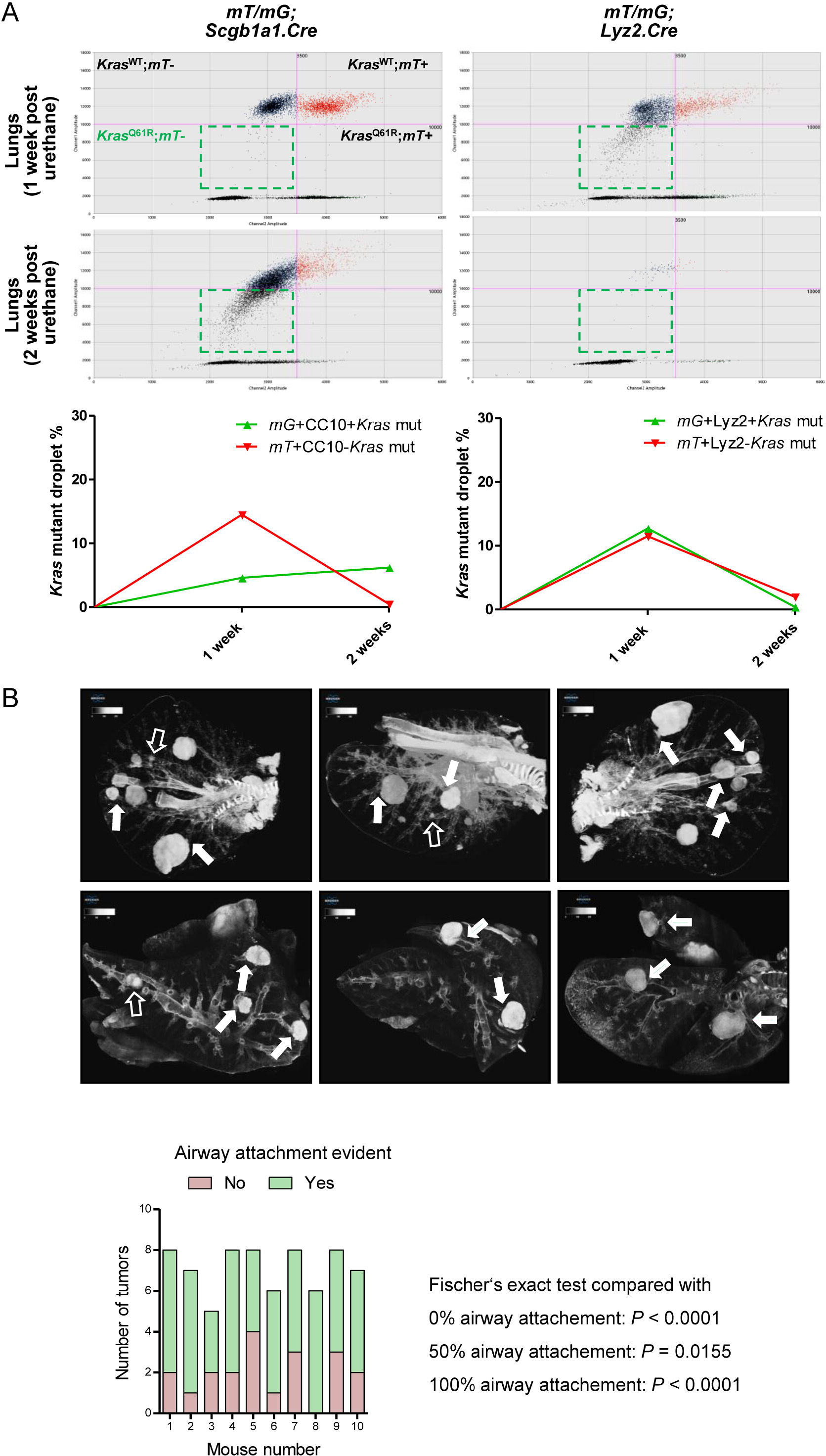
Airway cells contribute to the initiation of chemical-induced LUAD. **A,** Amplitude graphs (top two rows) and relative quantifications (bottom graphs) of digital droplet PCR (ddPCR) with probes targeting the *mT* and the *Kras*^Q61R^ sequences performed on DNA samples extracted from lungs of *mT/mG;Scgb1a1.Cre* and *mTmG;Lyz2.Cre FVB* mice treated with urethane and harvested one and two weeks post carcinogen insult. Note that only the *mG*+;*Kras*^Q61R^ cells (contained in the green dashed areas of the amplitude graphs) of the *mT/mG;Scgb1a1.Cre* mice (i.e. the CC10+ AEC with *Kras*^Q61R^ mutations) maintained the *Kras*^Q61R^mutation. **B,** Representative high resolution micro-computed tomography (micro-CT) lung sections (top row) and three-dimensional reconstruction images (bottom row) of urethane treated *FVB* mice six months into treatment (*n* = 10). Note the close association of most LUAD with the airways, as lung tumors seem to stem from the airways (arrows) or even to be contained in them (open arrows). Graph shows lung tumor attachment to bronchial structures as evident by three-dimensional reconstruction of ten lungs.

### Dissemination of airway signatures across the tumor-initiated lung

Since airborne carcinogens act globally on the respiratory field, we examined non-neoplastic alveolar areas of carcinogen-treated *mT/mG;Scgb1a1.Cre*, *mT/mG;Sftpc.Cre*, and *mT/mG;Lyz2.Cre* mice. Dissemination of the airway transcriptomic signature was evident by the markedly increased *mG*+ cell numbers in the alveoli of carcinogen-treated *mT/mG;Scgb1a1.Cre* mice compared with saline-treated controls (Figure 3A and Figure S4A and S4B). Immunostaining revealed that these *Scgb1a1*+ marked cells were CCSP+TUBA1A-SFTPC- when located near airways and CCSP-TUBA1A-SFTPC+ in alveoli and tumors (Figure 3B-3D and Figure S4C). Expansion of *Scgb1a1*+ marked cells after urethane treatment was also documented using bioluminescent imaging of double heterozygote offspring of *R26.Luc* [26] intercrosses with *Scgb1a1.Cre* mice, yielding a strain featuring light emission from *Scgb1a1*+ cells (Figure S4D and S4E). In addition, co-staining of human LUAD [27] for SFTPC, CCSP, and KRT5 showed co-localization of SFTPC with KRT5 but not with CCSP (Figure 3E-3G). These results indicate that dynamic changes in alveolar cell composition and/or gene expression occur globally during field cancerization by tobacco carcinogens. Altogether, the findings suggest that some alveolar cells are recycled by *Scgb1a1+* cells or transiently express CCSP during carcinogenesis, resulting in expansion of the *Scgb1a1*+ signature across the lungs. Moreover, that human and murine LUAD carry mixed epithelial signatures although their location and protein expression suggests an alveolar origin [18, 28–30].

**Figure 3.**
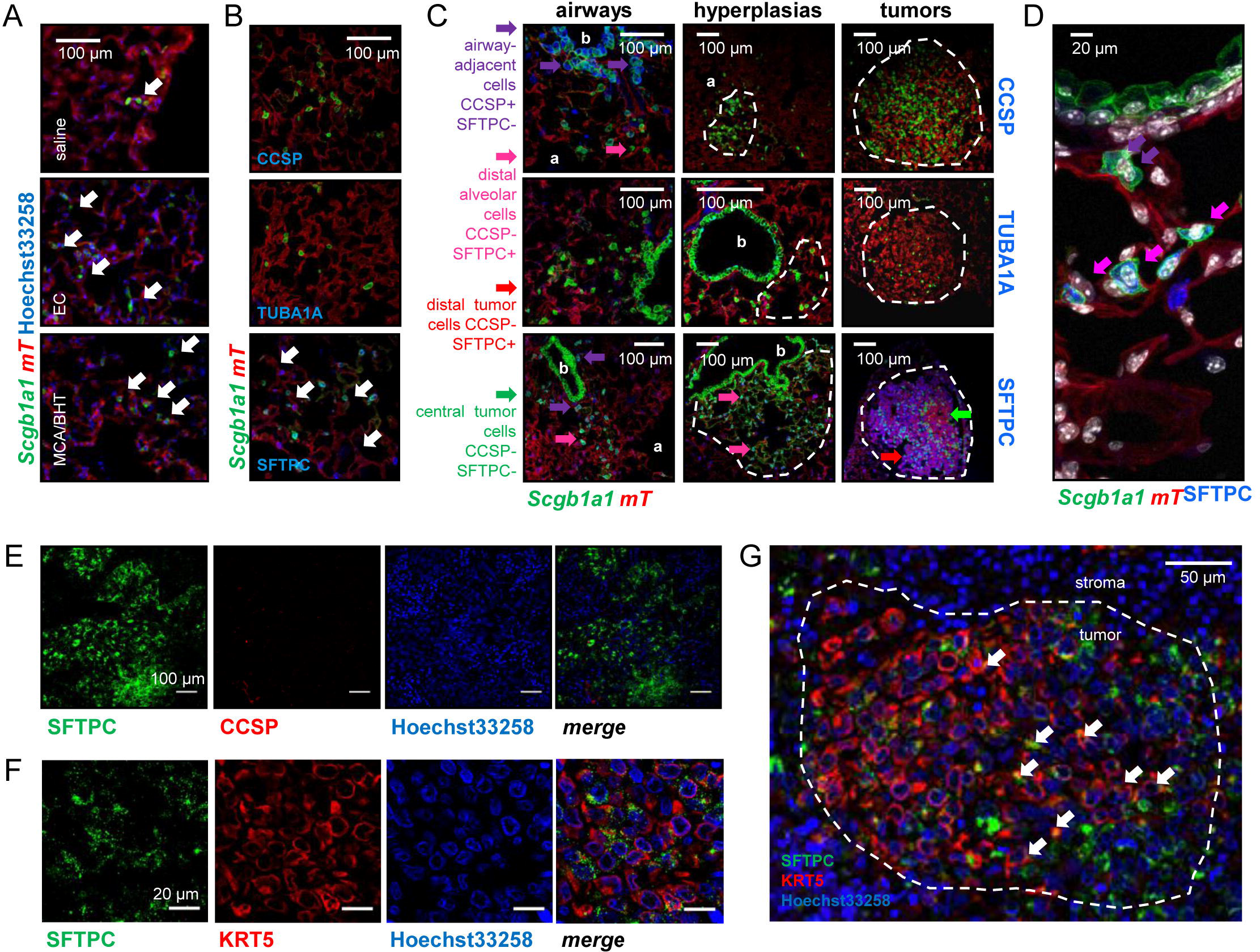
Expansion of airway-marked cells across the carcinogen-permutated respiratory field. **A,** Non-neoplastic alveolar regions of saline-, urethane-, and MCA/BHT-treated *C57BL/6 mT/mG;Scgb1a1.Cre* mice at six months into treatment (*n* = 10, 22, and 8/group, respectively). Note increased alveolar-located *mG*+ cell numbers in carcinogen-treated mice (arrows). **B,** Non-neoplastic distal lungs of urethane-treated *mT/mG;Scgb1a1.Cre* mice at six months into treatment (*n* = 22) stained for lineage markers. Note the alveolar *mG*+CCSP-TUBA1A-SFTPC+ cells (arrows). **C** and **D,** Juxtabronchial regions, alveolar hyperplasias and tumors (dashed lines) of lungs from urethane-treated *mT/mG;Scgb1a1.Cre* mice at six months into treatment (*n* = 22) stained for lineage markers. Arrows and legend indicate different phenotypes of extrabronchial *mG*+ cells. a, alveoli; b, bronchi. **E** and **F,** Co-staining of human lung adenocarcinomas (LUAD) for SFTPC and either CCSP (E; *n* = 10) or KRT5 (F; *n* = 10) shows absence of CCSP expression and significant co-localization of SFTPC and KRT5 in a subset of LUAD cells. **G,** Merged high-power image of SFTPC and KRT5 co-staining of human LUAD (dashed outline) from F. Arrows indicate SFTPC and KRT5 co-localization. Five non-overlapping fields/sample were examined. *mG*, membranous green fluorescent protein fluorophore; *mT*, membranous tomato fluorophore; CCSP, Clara cell secretory protein; TUBA1A, acetylated α-tubulin; KRT5, keratin 5; SFTPC, surfactant protein C; LYZ2, lysozyme 2.

### *Scgb1a1*+ marked cells in lung injury and repair

We next examined the dynamics of epithelial signatures during aging, injury, and repair. While *mG*+ cell abundance in the lungs of aging *mT/mG;Sftpc.Cre* and *mT/mG;Lyz2.Cre* mice did not change, *Scgb1a1*+ marked cells progressively increased in the alveoli of aging *mT/mG;Scgb1a1.Cre* mice, and these cells expressed SFTPC (Figure 4A). Bleomycin treatment, which depletes ATII cells [31], accelerated the accumulation of *Scgb1a1*+ marked cells in the alveoli and in urethane-triggered LUAD (Figure 4B and Figure S5A and S5B). Alveolar *Scgb1a1*+ cells also increased in response to perinatal hyperoxia that damages forming alveoli, as well as to naphthalene treatment that kills AEC (Figure 4C and 4D) [31]. However, no SFTPC-expressing or *Lyz2*+ marked AEC were found in naphthalene-treated *mT/mG;Scgb1a1.Cre* or *mT/mG;Lyz2.Cre* mice, respectively (Figure 4D and Figure S5C and S5D). Hence *de novo Scgb1a1*+ signatures appear in the alveolar regions of the aging and injured adult mouse lung, a fact that can be explained by peripheral lung migration of AEC or, as suggested by a previous study [32], by transient up-regulation of CCSP expression by regenerating alveolar cells. In addition, ATII-restricted transcriptomic signatures are not observed in the airways after injury, in line with previous work [14].

**Figure 4.**
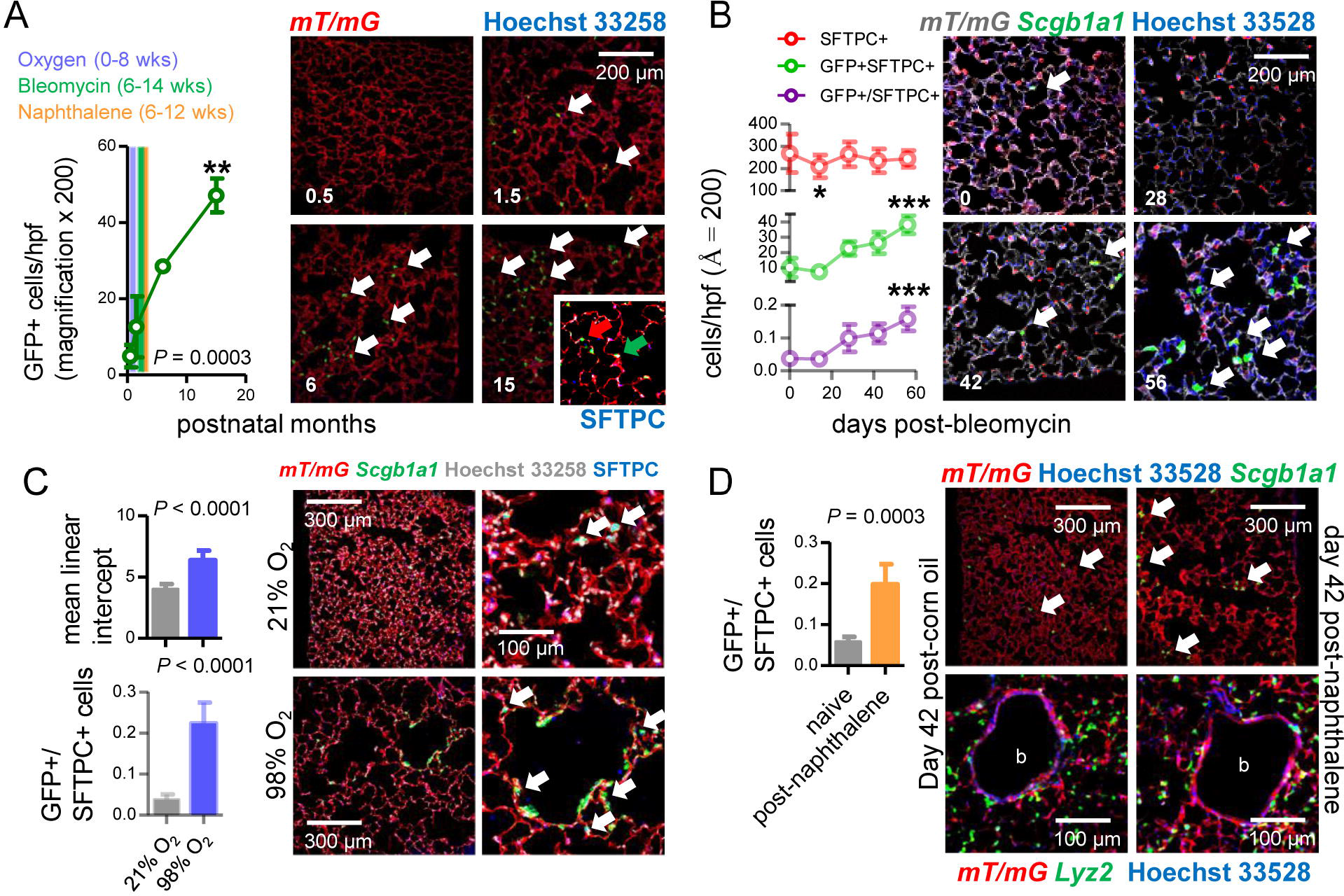
Airway-marked cells in alveolar repair. **A,** Data summary (graph; *n* = 5 mice/time-point) and lung sections of aging *mT/mG;Scgb1a1.Cre* mice show increasing alveolar *mG*+ cell abundance with age (arrows). Arrows in insert: *mG*+SFTPC+ (green) and *mG*+SFTPC- (red) airway-marked cells in alveolus of 15-month-old *mT/mG;Scgb1a1.Cre* mouse. Color-coded graph areas and corresponding text: time-windows of lung injury experiments. **B,** Data summary (graph; *n* = 4 mice/time-point) and SFTPC-stained sections of *mT/mG;Scgb1a1.Cre* lungs show accelerated increase of alveolar *mG*+SFTPC+ cells after bleomycin treatment (arrows). **C,** Data summary (graph; *n* = 6 mice/group) and SFTPC-stained sections of *mT/mG;Scgb1a1.Cre* lungs at two months after perinatal exposure to 98% O_2_ show enlarged alveoli (evident by increased mean linear intercept) enriched in *mG*+SFTPC+ cells (arrows) compared with 21% O_2_. **D,** Data summary (graph; *n* = 5 mice/group) and sections (top images) of *mT/mG;Scgb1a1.Cre* lungs show enrichment of alveoli in *mG*+ cells post naphthalene treatment (arrows). Bottom images: sections of *mT/mG;Lyz2.Cre* lungs (*n* = 5 mice/group) at six weeks post-naphthalene show no bronchial (b) *mG*+ cells. *mG*, membranous green fluorescent protein fluorophore; *mT*, membranous tomato fluorophore; CCSP, Clara cell secretory protein; SFTPC, surfactant protein C; LYZ2, lysozyme 2. Measurements were from five non-overlapping fields/lung. Data are given as mean ± SD and one-way ANOVA (A and B) or unpaired Student’s t-test (C and D) *P* values. *, **, and ***: *P*< 0.05, *P*< 0.01, and *P*< 0.001, respectively, for comparison with time-point zero by Bonferroni post-tests.

### Airway and alveolar contributions to alveolar maintenance and adenocarcinoma in the adult lung

To test the role of *Scgb1a1*+, *Sftpc*+, and *Lyz2*+ marked cells in alveolar homeostasis and carcinogenesis, we ablated them by crossing *Scgb1a1.Cre*, *Sftpc.Cre*, and *Lyz2.Cre* mice to *Dta* mice expressing *Diphtheria* toxin in somatic cells upon *Cre*-mediated recombination [33]. Triple transgenic *mT/mG;Driver.Cre;Dta* intercrosses were also generated to evaluate ablation efficiency. As expected, *Sftpc.Cre;Dta* and *mT/mG;Sftpc.Cre;Dta* mice were fetal lethal (no double or triple heterozygote offspring was obtained by *n* > 3 intercrosses, > 10 litters, and > 60 off-springs for each genotype; *P*< 0.0001 by Fischer’s exact test). However, all other ablated mice survived till adulthood. *Scgb1a1*+ cell ablation was complete in *mT/mG;Scgb1a1.Cre;Dta* mice, while only *Lyz2*+ AMФ persisted in *mT/mG;Lyz2.Cre;Dta* mice, which were freshly recruited monocytes just starting to express Lyz2 (Figure S5E and S5F). Immunostaining revealed that the denuded airway epithelium of 12-week-old *mT/mG;Scgb1a1.Cre;Dta* mice contained few flat CCSP+SFTPC+LYZ2+ cells, while the apparently intact alveolar spaces of *mT/mG;Lyz2.Cre;Dta* mice harbored CCSP-SFTPC-LYZ2+ AMФ (Figure 5A). Remarkably, morphometric and functional analyses of 12-week-old *Dta* control, *Scgb1a1.Cre;Dta*, and *Lyz2.Cre;Dta* mice showed that *Lyz2.Cre;Dta* mice displayed normal airway caliper and mean linear intercept (measures of bronchial and alveolar structure), normal number of CD45+CD11b+ myeloid cells in bronchoalveolar lavage (measure of airspace inflammation), as well as normal airway resistance and static compliance (measures of bronchial and alveolar function) compared with *Dta* controls. However, *Scgb1a1.Cre;Dta* mice displayed widened airway caliper, enlarged alveoli, and inflammatory interalveolar septal destruction evident by increased mean linear intercept, bronchoalveolar lavage of CD45+CD11b+ cells, and static compliance (Figure 5B and 5C), mimicking human chronic obstructive pulmonary disease [1]. Finally, we exposed control and ablated mice to ten consecutive weekly urethane exposures. All mice survived six months into carcinogen treatment, and *Scgb1a1.Cre;Dta* and *Lyz2.Cre;Dta* mice were equally protected from LUAD development compared with controls (Figure 5D). Taken together, these results show that *Scgb1a1*+ marked cells maintain postnatal alveolar structure and function, and, together with *Lyz2*+ marked cells, are required for LUAD development.

**Figure 5.**
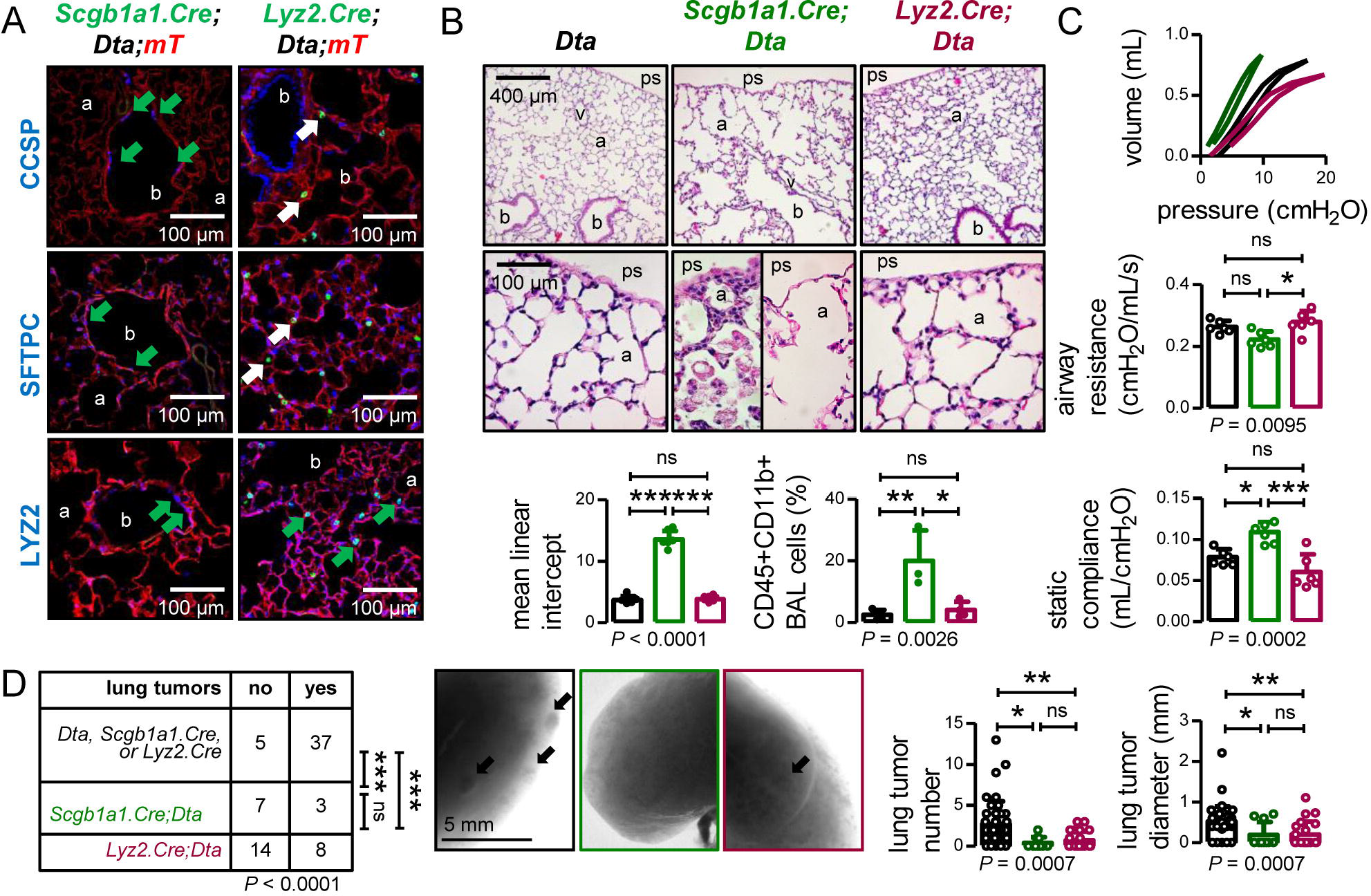
Airway-marked cells in alveolar maintenance and chemical-induced LUAD. **A,** Lineage marker-stained lung sections of 12-week-old *mT/mG;Scgb1a1.Cre;Dta* and *mT/mG;Lyz2.Cre;Dta* mice (*n* = 6/group) show increased bronchial and alveolar size and flat CCSP+SFTPC+LYZ2+ cells in the airways of *mT/mG;Scgb1a1.Cre;Dta* mice (green arrows), and CCSP-SFTPC- (white arrows) LYZ2+ (green arrows) alveolar macrophages in the airspaces of *mT/mG;Lyz2.Cre;Dta* mice. **B,** Hematoxylin and eosin-stained lung sections (top) and data summaries of mean linear intercept (bottom left, *n* = 6/group) and bronchoalveolar lavage (BAL) myeloid cells (bottom right, *n* = 7-10/group) from 12-week-old *Dta*, *Scgb1a1.Cre;Dta*, and *Lyz2.Cre;Dta* mice. **C,** Pressure-volume curves (top) and data summaries of airway resistance and static compliance from 12-week-old *Dta*, *Scgb1a1.Cre;Dta*, and *Lyz2.Cre;Dta* mice (*n* = 6/group). Note decreased airway resistance indicative of weakened airway walls and increased static compliance indicative of interalveolar septal destruction. **D,** Incidence (table), images (photographs), and data summaries of number and diameter of lung tumors of control, *Scgb1a1.Cre;Dta*, and *Lyz2.Cre;Dta* mice (*n* is given in table) at six months into treatment with urethane started at six weeks of age. a, alveoli; b, bronchi; ps, pleural space; v, vessel. *mG*, membranous green fluorescent protein fluorophore; *mT*, membranous tomato fluorophore; CCSP, Clara cell secretory protein; SFTPC, surfactant protein C; LYZ2, lysozyme 2. Five non-overlapping fields/sample were examined. Data are given as mean ± SD and one-way ANOVA (graphs) or χ^2^-test (table) *P* values. ns, *, **, and ***: *P*> 0.05, *P*< 0.05, *P*< 0.01, and *P*< 0.001, respectively, for the indicated comparisons by Bonferroni post-tests (graphs) or Fischer’s exact tests (table).

### Enrichment of airway and alveolar signatures in experimental and human lung adenocarcinoma

We subsequently cross-examined the transcriptomes of LUAD cell lines isolated from urethane-induced lung tumors [34, 35] and of their originating murine lungs with the gene expression profiles of murine AEC isolated from tracheal explants, of murine ATII cells [36], and of murine bone-marrow-derived macrophages (BMDM). The AEC signature was significantly enriched in LUAD cells compared with the ATII and BMDM signatures and with whole lungs (Figure 6A and 6B and Figure S6A). LUAD cell lines lost expression of several epithelial markers compared with their native lungs, but displayed markedly up-regulated expression of LUAD markers (i.e., *Krt18* and *Krt20*), of EGF receptor ligands (*Areg* and *Ereg*), and of the *Myc* oncogene (Figure S6B-S6E). Similar analyses of the transcriptomes of human LUAD and corresponding healthy lungs [37] and the profiles of primary human AEC, ATII cells, and AMΦ [38–40], also disclosed that the AEC (but not the ATII and AMΦ) signature was significantly enriched in LUAD compared with healthy lung tissue (Figure 6C and 6D). Gene set enrichment analyses showed that while AEC, ATII, and BMDM signatures were significantly enriched in murine lungs, the AEC signature predominated over ATII and BMDM signatures in LUAD cells. In addition, all human AEC, ATII and AMΦ signatures were significantly enriched in human LUAD compared with healthy lungs (Figure 6E and 6F and Figure S7). These results were plausible by the early nature of the human surgical specimens examined compared with our murine cell lines that represent advanced metastatic tumor-initiating cells, and collectively indicated the presence of an anticipated alveolar, but also an unexpected airway epithelial transcriptomic signature in tobacco carcinogen-induced LUAD.

**Figure 6.**
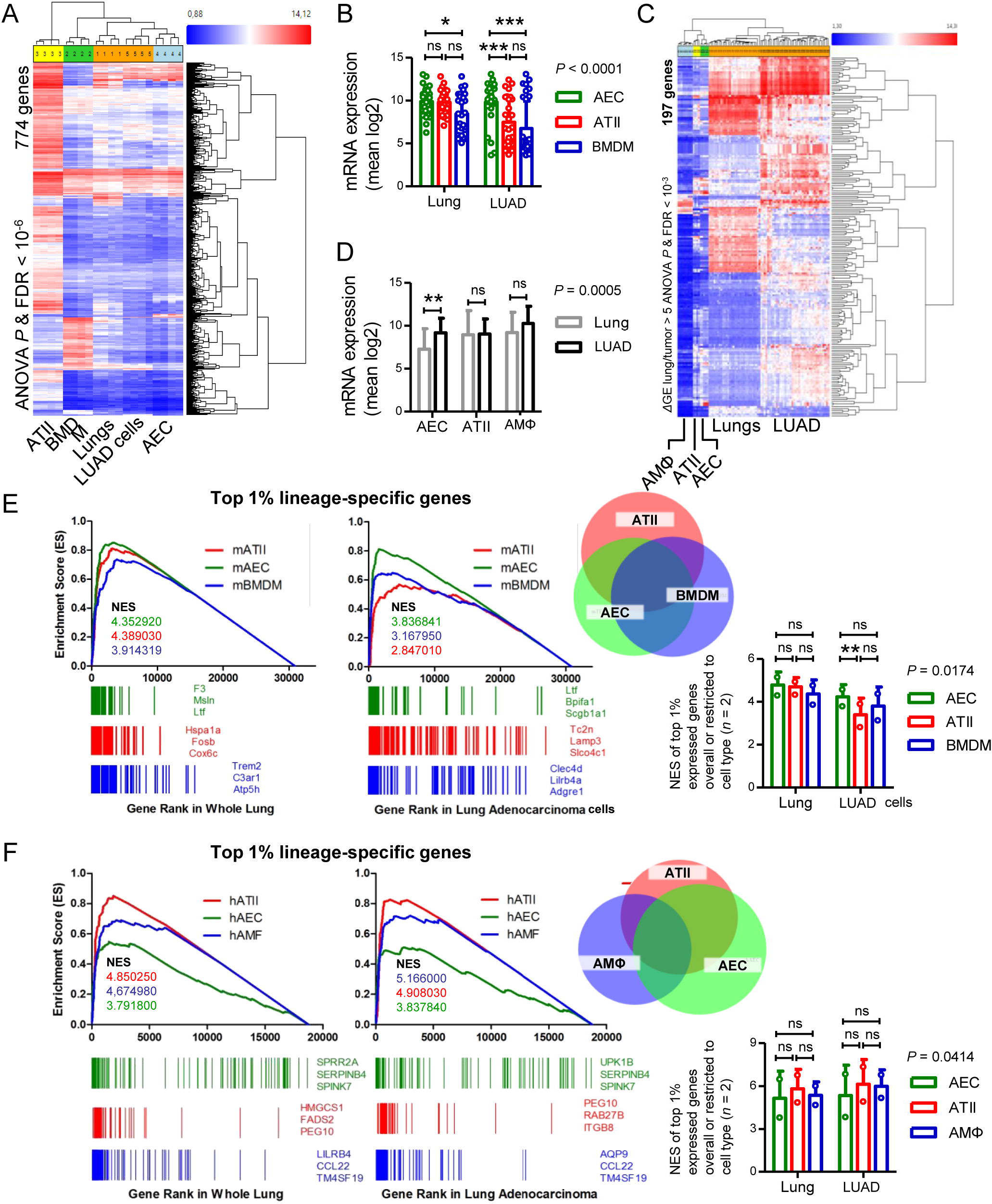
Airway and alveolar signatures in murine and human lung adenocarcinoma (LUAD). **A** and **B,** RNA of mouse urethane-induced LUAD cell lines, lungs obtained pre- and one week post-urethane treatment (*n* = 2 each), airway epithelial cells (AEC), alveolar type II cells [ATII; data from [36]], and bone marrow-derived macrophages (BMDM) was examined by Affymetrix Mouse Gene ST2.0 microarrays (*n* = 4/group). **A,** Heat map of genes significantly differentially expressed (overall ANOVA and FDR *P*< 10^−6^) shows accurate hierarchical clustering. **B,** Expression of the 30 top-represented transcripts of AEC, ATII, and BMDM in lungs and LUAD cells. **C** and **D,** RNA of human LUAD (*n* = 40), never-smoker lung tissue (*n* = 30), primary AEC (*n* = 5), primary ATII (*n* = 4), and alveolar macrophages (AMФ; *n* = 9) analyzed by Affymetrix Human Gene ST1.0 microarrays was cross-examined [data from [37–40]]. **C,** Heat map of genes significantly differentially expressed (*∆*GE > 5-fold) between LUAD and lung (ANOVA and FDR *P*< 10^−3^) shows accurate hierarchical clustering. **D,** Mean expression levels of the 30 top-represented transcripts of human AEC, ATII, and AMФ in lungs and LUAD. **E** and **F,** Gene set enrichment analysis, including normalized enrichment scores (NES), of mouse (E) and human (F) AEC, ATII, and BMDM/AMФ signatures (defined as the top 1% expressed genes overall or exclusive to the cell type) in mouse and human LUAD transcriptomes shows significant enrichment of the AEC (but not the ATII and BMDM/AMФ) signature compared with lung (nominal *P*< 0.0001 for all, family-wise error rates FWER < 0.01). Gene symbols indicate the top 3 lagging genes from each signature and shows loss of *Scgb1a1* by LUAD. Data are given as mean ± SD and two-way ANOVA *P* values. ns, *, **, and ***: *P*> 0.05, *P*< 0.05, *P*< 0.01, and *P*< 0.001 for the indicated comparisons by Bonferroni post-tests. ANOVA, analysis of variance; FDR, false discovery rate.

## DISCUSSION

We characterized the dynamics of respiratory epithelial signatures in the postnatal mouse lung during aging and after challenge with noxious and carcinogenic insults. The contribution of airway and alveolar signatures to chemical-induced LUAD of mice and men is described for the first time (Figure 7A). Although the peripheral location and molecular phenotype of these LUAD suggest an alveolar origin, we show here that both airway and alveolar-programmed cells are found in chemical-induced LUAD and that, in fact, AEC may play a more prominent role during the initial steps of chemical lung carcinogenesis. Furthermore, both airway and alveolar cells are implicated in postnatal alveolar maintenance during aging and recovery from injury. Our analyses facilitate insights into the dynamics of epithelial signatures in the postnatal lung (Figure 7B) and indicate that both airway and alveolar-restricted transcriptomic programs are essential for its sustained structural and functional integrity. Finally, urethane-induced mouse LUAD cells and human LUAD are shown to bare in their transcriptome markings of multiple pulmonary epithelial lineages including highly enriched airway signatures, rendering our findings plausible in both experimental murine and human LUAD.

**Figure 7.**
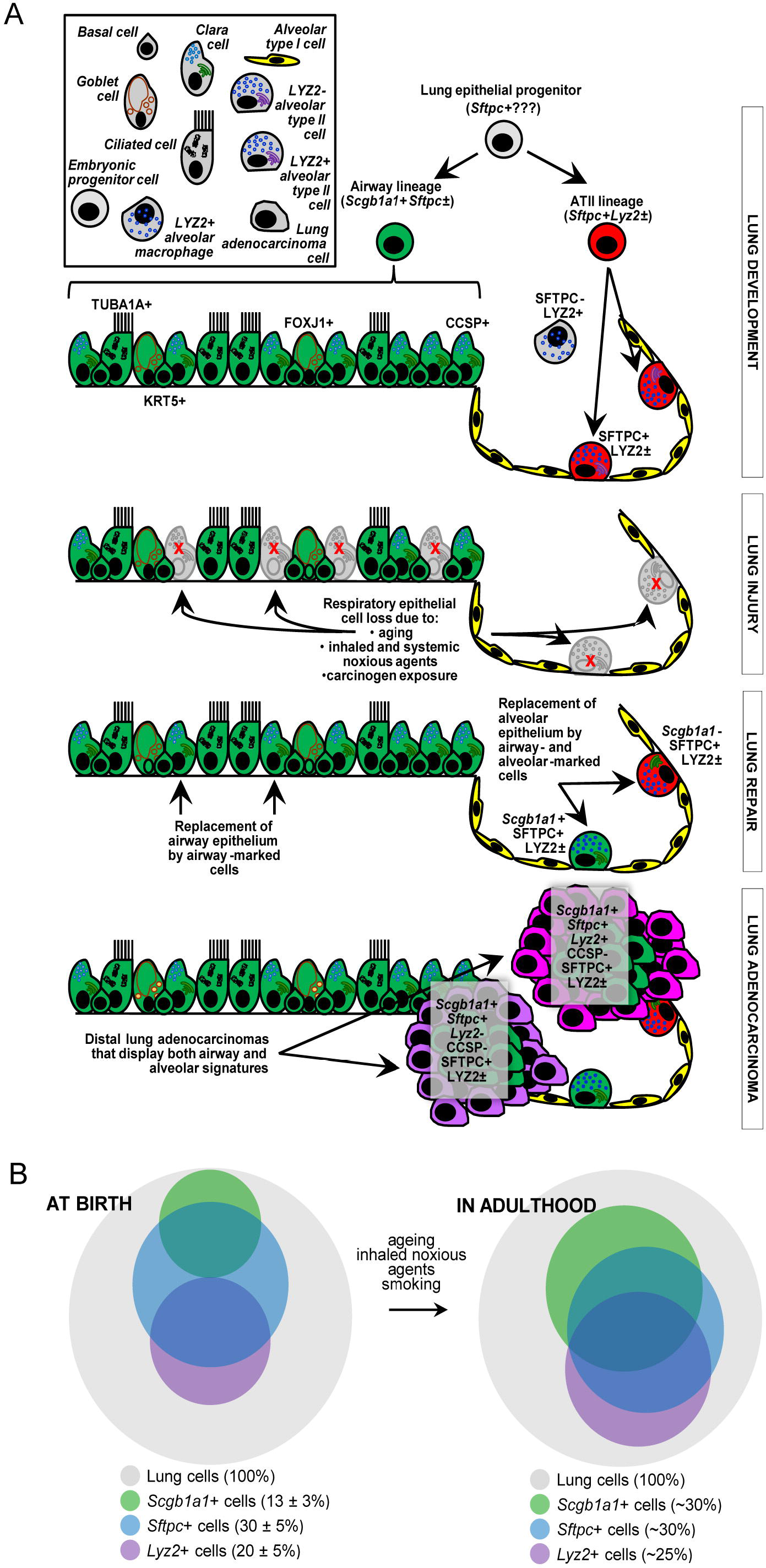
**A,** Proposed role of airway-marked cells in distal lung maintenance and adenocarcinoma. Our evidence supports the existence of distinct developmental ancestries for airway and alveolar type II (ATII) cells, notwithstanding their common descent from an early (possibly *Sftpc*+) lung epithelial progenitor. The developmental airway lineage (*Scgb1a1*+*Sftpc*±; green) gives rise to all types of airway cells, including club or Clara, ciliated, goblet, basal, and other cells, while the developmental ATII lineage (*Sftpc*+*Lyz2*±; red) gives rise to ATII cells formed before birth. These lineages appear to be relatively segregated in the growing unaffected lung of the mouse till the age of six weeks, which roughly corresponds to a human age of six years, where cellular proliferation in the human lungs ceases. Thereafter, and likely due to the continuous bombardment of the lungs by inhaled noxious particles and substances during normal respiration, gradual expansion of *Scgb1a1*+*Sftpc*± marked cells ensues. Upon lung injury, this process is markedly accelerated. Similarly, during carcinogenesis caused by chemical carcinogens of tobacco smoke, we show how *Scgb1a1*+*Sftpc*± marked cells expand and are ubiquitously present in even peripheral lung adenocarcinomas. **B,** Proposed neonatal proportions and postnatal dynamics of pulmonary epithelial signatures during adulthood. Estimated proportions of lineage-marked cells at birth, based on flow cytometry and co-localization of proteinaceous and genetic cell marking (Figure 1A and Figure S1). Lung lineages appear to be relatively segregated in the growing lung till the age of full lung development (six weeks in mice and 6-8 years in humans) or till lung injury ensues, whichever comes first. Schematic of postnatal redistribution of marked cells in the adult lung, based on the findings of the present work and a previous report [14]. Upon injury to airway and/or alveolar cells, during multi-stage field carcinogenesis, or even during unchallenged aging, *Scgb1a1*+ marked cells appear in the distal alveolar regions, thereby maintaining lung structure and function. Bubble size indicates relative marked cell abundance. CCSP, Clara cell secretory protein; FOXJ1, forkhead box J1; KRT5, keratin 5; LYZ2, lysozyme 2; SFTPC, surfactant protein C; TUB1A1, acetylated α-tubulin.

This study addresses the cellular and molecular signatures of chemical-induced LUAD. Lung tumors induced in two different mouse strains by two different chemical regimens contained in tobacco smoke, urethane and MCA/BHT, are shown to contain multiple epithelial markings and signatures. This is important because human LUAD is inflicted by chronic exposure to tobacco smoke and other environmental exposures [41]. As such, the mutation profile of the human disease is more closely paralleled by chemical-induced murine lung tumors compared with lung cancers triggered by transgenic expression of *Kras*^G12C^ or *Kras*^G12D^ in the respiratory epithelium [15]. Although the latter transgenic tumors have been extensively studied [9–14], chemical-induced LUAD have not been investigated. In both of the mouse models that we used, even in the *FVB* one-hit model involving a single dose of carcinogen administration, all developed LUAD contained the *Scgb1a1*+ genetic marking, in contrast with the *Lyz2*+ genetic marking which was dispensable for LUAD formation. These observations could support a multi-stage course of events in chemical carcinogenesis, involving at some point an airway-specific transcriptomic signature. In fact, the prevalence of a different *Kras* mutation in urethane-induced tumors (*Kras*^Q61R^ compared to *Kras*^G12^ mutations in the transgenic mouse models) has led to the suggestion that chemical carcinogens introduce *Kras* mutations in a different population of tumor-initiating cells than the mouse models of genetic activation of *Kras* [15]. Our findings of *Scgb1a1*+ AEC being more sensitive than *Lyz2*+ ATII cells to*Kras*^Q61R^ mutations during the initiation steps of urethane-induced lung carcinogenesis further supports this notion. The findings of chemical-triggered LUAD, as well as of their precursor hyperplastic lesions, bearing *Scgb1a1*+, *Sftpc*+, and *Lyz2*+ markings, implies that they can originate from: i) AEC that colonize the distal lung during carcinogenesis thereby activating obligate (*Sftpc*+) and dispensable (*Lyz2*+) alveolar transcriptomic programs; ii) alveolar cells that transit through an obligate *Scgb1a1*+ and a dispensable *Lyz2*+ stage during the process; or iii) multipotent progenitors that express multiple epithelial signatures, such as those found during pulmonary embryogenesis, in human LUAD, and in other chronic lung diseases [40–44]. However, in our view, the propensity of airway cells to survive *Kras*^Q61R^ mutations during the early initiation steps of urethane-induced lung carcinogenesis, and the close airway association of lung tumors revealed by our high resolution μCT analysis support a bronchial origin of these tumors, in line with recent evidence of tobacco smoke inducing epigenetic changes that sensitize human airway epithelial cells to a single *KRAS* mutation [45]. Along these lines, the split phenotype of chemical-induced lung tumors of *mT/mG;Lyz2.Cre* mice indicates that *Lyz2*-expressing ATII cells can be dispensable for carcinogen-triggered LUAD development, as opposed to what has been previously shown for genetically-triggered LUAD [14].

Our approach focuses on integral assessment of the lung epithelial transcriptomic signatures participating in adult lung changes in response to aging, injury, and carcinogenesis. The identification of transcriptomic programs and signatures that are activated during lung repair and carcinogenesis and that team up with oncogenic signaling in a non-oncogenic addictive fashion is of great importance for LUAD biology and is likely to lead to therapeutic innovations [46]. To this end, it was recently shown that insertions and deletions in lineage-restricted genes occur in human LUAD [7]. Moreover, integrin β_3_ and TANK-binding kinase 1 partner with oncogenic *KRAS* signaling to mediate cancer stemness and drug resistance [5, 6]. Along these lines, our findings of both airway and alveolar transcriptomes being involved in lung maintenance, repair, and carcinogenesis is of great importance, as it implies that these programs facilitate the survival and proliferation of lung stem cells with regenerative potential and of mutated cells with malignant potential and are thus therapeutic targets. The perpetual cell marking approach adopted was preferential over available pulsed lineage tracing models because of the unprecedented accuracy of our *Scgb1a1.Cre* strain in exclusively and completely marking airway epithelial cells at the conclusion of development, allowing tracking of subsequent changes in adulthood.

In conclusion, airway cells and transcriptomic programs contribute to alveolar maintenance and LUAD. Since defective epithelial repair underlies the pathogenesis of chronic lung diseases and since abundantly transcribed genes are central to the mutational processes that cause cancer, this finding is of potential therapeutic importance for chronic pulmonary diseases and lung cancer.

## ACKNOWLEDGMENTS

The authors thank the University of Patras Centre for Animal Models of Disease and Advanced Light Microscopy Facility for experimental support. M.A. and F.R. also wish to thank the InfrafrontierGR infrastructure (co-financed by the ERDF and NSRF 2007-2013) for the excellent microCT facilities.

## AUTHOR CONTRIBUTIONS

MS designed and carried out experiments, analyzed the data, and wrote the paper draft; IL, YC, DEZ, MV, NIK, ADG, VAr, KAMA, LVK, DT, VK, SGZ and IG carried out experiments and analyzed the data; MAP and ASL performed digital droplet PCR; MA and FR performed micro-CT analysis; DEW performed gene set enrichment analyses; AM provided critical experimental and intellectual input; VAi and RS provided critical analytical tools, designed experiments, analyzed the data, and edited the paper; GTS conceived the main idea, obtained funding and supervised the study, provided analytical tools, designed experiments, analyzed the data, wrote the paper, and is the guarantor of the study’s integrity. All authors reviewed the manuscript and concur with the submitted version.

## METHODS

### Experimental mice

*C57BL/6*J (*C57BL/6*; #000664), *FVB/NJ* (#001800), *B6.129(Cg)-Gt(ROSA)26Sor*^*tm4(ACTB-tdTomato,-EGFP)Luo*^/*J* [*mT/mG*; #007676; [19]], *FVB.129S6(B6)-Gt(ROSA)26Sor*^*tm1(Luc)Kael*^/*J* [*R26.Luc*; #005125; [26]], *B6.129P2-Gt(ROSA)26Sor*^*tm1(DTA)Lky*^/*J* [*Dta*; #009669; [33]], *B6.129P2-Lyz2*^*tm1(cre)Ifo*^/*J* [*Lyz2.Cre*; #004781; [14]], *B6.Cg-Tg(Sox2-cre)1Amc/J* [*Sox2.Cre*; #008454; [22]], *B6.Cg-Tg(Vav1-icre)A2Kio/J* [*Vav.Cre*; #008610; [23]], and *B6.Cg-Tg(Nes-cre)1Kln/J* [*Nes.Cre*; #003771; [24]] mice were from Jackson Laboratories (Bar Harbor, MN). *B6;CBA-Tg(Scgb1a1-cre)1Vart/Flmg* (*Scgb1a1.Cre*; European Mouse Mutant Archive #EM:04965) mice were developed by our group [20] and *Tg(Sftpc-cre)1Blh* (*Sftpc.Cre*; Mouse Genome Informatics #MGI:3574949) mice were donated by their founder [21]. Mice were bred >F12 to the *FVB* background at the University of Patras Center for Animal Models of Disease.

### Mouse models of lung adenocarcinoma

Six-week-old mice on the *C57BL/6* background received ten consecutive weekly intraperitoneal urethane injections (1 g/Kg in 100 μL saline) and were sacrificed 6-7 months after the first injection, or four consecutive weekly intraperitoneal MCA (15 mg/Kg in 100 μL saline) followed by eight consecutive weekly intraperitoneal BHT injections (200 mg/Kg in 100 μL corn oil) and were sacrificed 6-7 months after the first injection. Six-week-old mice on the *FVB* background received one intraperitoneal urethane injection (1 g/Kg in 100 μL saline) and were sacrificed 6-7 months later [16–18].

### Mouse models of lung injury

Six-week-old mice (*C57BL/6* background) received intratracheal bleomycin A2 (0.08 units in 50 μL saline) or intraperitoneal naphthalene (250 mg/Kg in 100 μL corn oil) [31, 32]. In addition, preterm mothers of the *C57BL/6* background and their offspring were exposed to room air (21% oxygen; control) or 98% oxygen for two days before and four days after birth [47].

### Mouse model of perinatal hyperoxic lung injury

For induction of perinatal hyperoxic lung injury, preterm mothers of the *C57BL/6* background and their offspring were exposed to room air (21% oxygen; control) or 98% oxygen for two days before and four days after birth, when oxygen-exposed pups were returned to room air, as described previously [47]. Oxygen levels were continuously monitored with an oxygen sensor. The gas stream was humidified to 40–70% by a deionized water-jacketed Nafion membrane tubing and delivered through a 0.22 μm filter before passage into a sealed Lexan polycarbonate chamber measuring 40 × 25 × 25 cm and accommodating 25 L gas at a flow rate of 5 L/min, resulting in complete gas exchange every 5 min. Mothers were cycled between litters on 21% and 98% oxygen every 24 hours to prevent oxygen toxicity and to control for nutritional support of the pups. After perinatal hyperoxia, mice remained at room air till sacrificed at eight weeks of age.

### Urethane-induced lung adenocarcinoma cell lines

Lung tumors were dissected from surrounding healthy lung parenchyma under sterile conditions, minced into 1-mm pieces, and cultured at 37°C in 5% CO2-95% air using DMEM 10% FBS, 2 mM L-glutamine, 1 mM pyruvate, 100 U/mL penicillin, and 100 U/mL streptomycin. All cell lines were immortal and indefinitely phenotypically stable over > 18 months and/or 60 passages, and were tumorigenic and metastatic in *C57BL/6* mice [34, 35].

### Human lung adenocarcinomas

Ten archival formalin-fixed, paraffin-embedded tissue samples of patients with LUAD that underwent surgical resection with curative intent between 2001 and 2008 at the University Hospital of Patras were retrospectively enrolled [27].

### Statistical analysis

Sample size was calculated using power analysis on G*power, assuming *α* = 0.05, *β* = 0.05, and effect size *d* = 1.5. No data were excluded from analyses. Animals were allocated to treatments by alternation and transgenic animals were enrolled case-control-wise. Data were collected by at least two blinded investigators from samples coded by non-blinded investigators. All data were normally distributed by Kolmogorov-Smirnov test, are given as mean ± SD, and sample size (*n*) always refers to biological and not technical replicates. Differences in frequency were examined by Fischer’s exact and χ^2^ tests and in means by t-test or one-way ANOVA with Bonferroni post-tests. Changes over time and interaction between two variables were examined by two-way ANOVA with Bonferroni post-tests. All probability (*P*) values are two-tailed and were considered significant when *P*<0.05. All analyses and plots were done on Prism v5.0 (GraphPad, La Jolla, CA).

## SUPPLEMENTAL INFORMATION

Supplemental information includes supplemental methods and seven figures and can be found with this article online.

