## Supplemental Methods for "Dual airway and alveolar contributions to adult lung homeostasis and carcinogenesis"

#### Study approval

All mice were bred at the Center for Animal Models of Disease of the University of Patras. Experiments were designed and approved *a priori* by the Veterinary Administration of the Prefecture of Western Greece (approval numbers 3741/16.11.2010, 60291/3035/19.03.2012, and 118018/578/30.04.2014) and were conducted according to Directive 2010/63/EU (http://eur-lex.europa.eu/legal-content/EN/TXT/?qid=1486710385917&uri=CELEX:32010L0063). Male and female experimental mice were sex-, weight (20-25 g)-, and age (6-12 week)-matched. *N* = 588 experimental and 165 breeder mice were used for this report. Sample size was calculated using power analysis on G*power. Experiments were randomized across different cages and mouse lungs were always examined by two blinded researchers. Sample numbers are included in the figures and figure legends.

#### Reagents

Urethane, ethyl carbamate, EC, CAS# 51-79-6; 3-methylcholanthrene, 3-methyl-1,2-dyhydrobenzo[j]aceanthrylene, MCA, CAS# 56-49-5; butylated hydroxytoluene, 2,6-Di-tert-butyl-4-methylphenol, BHT, CAS# 128-37-0; naphthalene, CAS# 91-20-3, and Hoechst33258 nuclear dye (CAS# 23491-45-4), were from Sigma-Aldrich (St. Louis, MO). Bleomycin A2, ((3-{[(2'-{(5S,8S,9S,10R,13S)-15-{6-amino-2- [(1S)-3-amino-1-{[(2S)-2,3-diamino-3-oxopropyl]amino}-3-oxopropyl] -5-methylpyrimidin-4-yl}-13-[{[(2R,3S,4S,5S,6S)-3-{[(2R,3S,4S,5R,6R)-4-(carbamoyloxy)-3,5-dihydroxy-6- (hydroxymethyl) tetrahydro-2H-pyran-2-yl]oxy} -4,5-dihydroxy-6-(hydroxymethyl) tetrahydro-2H-pyran-2-yl]oxy} (1H-imidazol-5-yl)methyl]-9-hydroxy-5-[(1R)-1-hydroxyethyl]-8,10-dimethyl-4,7,12,15-tetraoxo-3,6,11,14-tetraazapentadec-1-yl}-2,4'-bi-1,3-thiazol-4-

yl)carbonyl]amino}propyl)(dimethyl)sulfonium; CAS #9041-93-4, was from Calbiochem (Darmstadt, Germany). D-Luciferin potassium salt, (4S)-2-(6-hydroxy-1,3-benzothiazol-2-yl)-4,5-dihydrothiazole-4-carboxylic acid, CAS #2591-17-5, was from Gold Biotechnology (St. Louis, MO).

**Micro-computed tomography**

Urethane or saline treated *FVB* mice were sacrificed six months post urethane/saline injection. Lungs were inflated and fixed with 10% formalin overnight. They were then dehydrated and chemically dried for micro-CT scanning using a method kindly provided by Jeroen Hostens, Bruker. Briefly, a gradient ethanol dehydration protocol (from 70-100%) was applied, followed by a 2h incubation in Hexamethyldisilazane (HMDS; Sigma, St. Louis, MO) and a 2h air-drying. The dehydrated lungs were then scanned in the SkyScan 1172 scanner from Bruker (Kontich, Belgium) at 41kV without filtration and with 5.94 μm voxel resolution (exposure: 440 msec). The X-ray projections were obtained at 0.35° intervals with a scanning angular rotation of 180° and two frames were averaged for each rotation under a mean of 10 frames per random movement. 3D reconstructions were performed using NRECON (Bruker, Kontich, Belgium) software. Regions of interest for the whole lung and peripheral lung tissue were defined in the CT analysis software (CTan; Bruker, Kontich, Belgium), thresholds applied to detect tissue from background, and a 3D volume rendering of the lungs were performed using the CTVox software (Bruker, Kontich, Belgium).

**Structural assessments in murine lungs**

Mouse lungs were recoded (blinded) by laboratory members not participating in these studies and were always examined by two independent blinded participants of this study. The results obtained by each investigator were compared, and lungs were re-evaluated if deviant by > 20%. Lungs and lung tumors were initially inspected macroscopically under a Stemi DV4 stereoscope equipped with a micrometric scale incorporated into one eyepiece and an AxiocamERc 5s camera (Zeiss, Jena, Germany) in trans-illumination mode, allowing for visualization of both superficial and deeply-located lung tumors [18]. Tumor location was charted and diameter (δ) was measured. Tumor number (multiplicity) per mouse was counted and mean tumor diameter per mouse was calculated as the average of individual diameters of all tumors found in a given mouse lung. Individual tumor volume was calculated as $\pi\delta^3/6$. Mean tumor volume per mouse was calculated as the average of individual volumes of all tumors found in a given mouse lung, and total lung tumor burden per mouse as their sum. Following macroscopic mapping of lung and lung tumor morphology, lungs of fluorescent reporter mice were imaged on a Leica MZ16F fluorescent stereomicroscope equipped with GFP and RFP filters and a DFC 300FX camera (Leica Microsystems, Heidelberg, Germany) in order to determine their macroscopic fluorescent pattern (Figures S1A, S2B and S3B). Lung volume was measured by saline immersion, and lungs were embedded in paraffin, randomly sampled by cutting 5 μm-thick lung sections ($n$ = 10/lung), mounted on glass slides, and stained with hematoxylin and eosin for morphometry and histologic typing of lung tumors. For this, a digital grid of 100 intersections of vertical lines (points) was superimposed on multiple digital images of all lung sections from lung tissue of a given mouse using Fiji academic freeware. Total lung tumor burden was determined by point counting of the ratio of the area occupied by neoplastic lesions versus total lung area and by extrapolating the average ratio per mouse to total lung volume [48]. The results of this stereologic approach were compared with the macroscopic method detailed above, and were scrutinized if deviant by > 20%. In a final step of analyses, point counting of tissue sections was redone, this time assigning points to three different neoplasia categories: airway and/or alveolar hyperplasias, adenomas, and adenocarcinomas, all present in our carcinogen-initiated lungs, according to our previous published experience [18]. Based on the results of these analyses, the absolute number and percentage of hyperplasias, adenomas, and adenocarcinomas was calculated. Point counting of the ratio of the area occupied by each distinct type of neoplastic lesions versus total lung area and extrapolating the average ratio per mouse to total lung volume was used to calculate the absolute volume of hyperplasias, adenomas, and adenocarcinomas in each lung and their contribution towards total lung tumor burden. To evaluate alveolar structure and size, we calculated mean linear intercept using randomly sampled hematoxylin and eosin-stained lung sections, as described elsewhere [48]. For this, a digital grid of twenty random horizontal lines was superimposed on multiple digital images of all lung sections from lung tissue of a given mouse using Fiji. Mean linear intercept was calculated by counting the intercepts of interalveolar septae with the lines and the formula: $\Sigma\{2 \times (\text{length of line/ number of intercepts})\}$/total number of lines. All quantifications were done by counting at least five random non-overlapping fields of view of at least ten sections per lung.

**Histology and molecular phenotyping**

For histology, lungs were inflated to 20 cm$H_2O$ pressure that provides for a lung volume equivalent to the resting volume of the lungs (a.k.a. functional residual capacity in humans) and enables precise histologic observations on airway and alveolar structure avoiding false interpretations resulting from the study of compressed or over-inflated lungs [48]. Subsequently, lungs were fixed with 10% formalin overnight and were embedded in paraffin. Five-μm-thick paraffin sections were then counterstained with hematoxylin and eosin (Sigma, St. Louis, MO) and mounted with Entellan new (Merck Millipore, Darmstadt, Germany). For immunofluorescence, lungs were inflated with a 2:1 mixture of 4% paraformaldehyde:Tissue-Tek (Sakura, Tokyo, Japan), fixed in 4% paraformaldehyde overnight at 4°C, cryoprotected with 30% sucrose, embedded in Tissue-Tek and stored at -80$^{o}$C. Ten-μm cryosections were then post-fixed in 4% paraformaldehyde for 10 min, treated with 0.3% Triton X-100 for 5 min, and incubated in blocking solution containing 10% fetal bovine serum (FBS), 3% bovine serum albumin (BSA), 0.1% polyoxyethylene (20) sorbitanmonolaurate (Tween 20) in 1x phosphate-buffered saline (PBS) for 1 h. Following labeling with the indicated primary antibodies overnight at 4°C, sections were incubated with fluorescent secondary antibodies, counterstained with Hoechst 33258 and mounted with Mowiol 4-88 (Calbiochem, Darmstadt, Germany). The following primary antibodies were used: rabbit anti-PCNA (1:3000, ab2426, Abcam, London, UK), rabbit anti-LYZ2 (marking ATII cells and alveolar macrophages, 1:50, ab108508, Abcam), rabbit anti-KRT5 (marking basal cells, 1:200, ab53121, Abcam), rabbit anti-SFTPC (marking ATII cells, 1:200, sc-13979, Santa Cruz, Dallas, TX), rabbit anti-CCSP (marking Clara or club cells, 1:200, sc-25555, Santa Cruz), goat anti-CCSP (marking Clara or club cells, 1:1000, sc-9772, Santa Cruz), mouse anti-acetylated α-tubulin (marking ciliated cells, 1:2000, T7451, Sigma-Aldrich, St. Lewis, MO), rabbit anti-SFTPC (marking ATII cells, 1:500, AB3786, Merck-Millipore, Burlington, MA), and mouse anti-KRT5 (marking basal cells, 1:200, MA5-17057, Thermo Fisher Scientific, Waltham, MA). Alexa Fluor donkey anti-rabbit 488 (A21206, Thermo Fisher Scientific), Alexa Fluor donkey anti-mouse 568 (ab175700, Abcam), Alexa Fluor donkey anti-goat 568 (A11057, Thermo Fisher Scientific), Alexa Fluor donkey anti-rabbit 647 (A31573, Thermo Fisher Scientific), and Alexa Fluor donkey anti-mouse 647 (A31571, Thermo Fisher Scientific) secondary antibodies were used at 1:500 dilution. For isotype control, the primary antibody was omitted. Bright-field images were captured with an AxioLab.A1 microscope connected to an AxioCamERc 5s camera (Zeiss, Jena, Germany) whereas fluorescent microscopy was carried out either on an Axio Observer D1 inverted fluorescent microscope (Zeiss, Jena, Germany) or a TCS SP5 confocal microscope (Leica Microsystems, Wetzlar, Germany) with 20x, 40x and 63x lenses. Digital images were processed with Fiji. All quantifications of cellular populations were obtained by counting at least five random non-overlapping bronchial-, alveolar-, hyperplasia-, or tumor-containing fields of view per section.

### Pulmonary function testing

Following anesthesia induced by intraperitoneal ketamine (100 mg/Kg) and xylazine (10 mL/Kg) and tracheostomy, mice were mechanically ventilated by a Flexivent rodent ventilator (Scireq, Montreal, Ontario, Canada). The whole procedure, described elsewhere [49], lasted 15 min. After a 3-min run-in period of ventilation with 21% oxygen, a tidal volume of 10 mL/Kg, a respiratory rate of 150 breaths/min, and a positive end-expiratory pressure of 3 $cmH_2O$, paralysis was induced using 8 mg/Kg intraperitoneal succinyl choline, and total respiratory system impedance was obtained by applying an 8-sec-long pseudorandom frequency oscillation (0.5-19.75 Hz) to the airway opening. Thirty seconds prior to initiation of measurements, lung volume history was once controlled by a 6-sec-long inflation to 30 cm $H_2O$ pressure. Measurements were repeated thrice at 60 sec intervals and were averaged. Data were fit into the constant phase model in order to fractionate total respiratory input impedance into airways resistance (Raw) and tissue damping and elastance coefficients. To obtain pressure-volume (PV) curves, the respiratory system was incrementally inflated and deflated to 40 mL/Kg total volume at seven steps each and airway pressures were recorded on each volume change. The slope of the linear portion of expiratory PV curves, which represents static compliance (Cst), a measure of airspace function, was calculated manually. Operators were blinded to animal genotype.

### Digital droplet PCR

*mT/mG* (n=4), *mT/mG;Scgb1a1.Cre* (n=4) and *mT/mG;Lyz2.Cre* (n=4) mice were injected with one intraperitoneal injection of urethane (1 g/Kg) and lungs were then harvested one (n=2) and two (n=2) weeks post urethane administration, homogenized and subjected to DNA extraction and purification using GenElute Mammalian Genomic DNA Minipreps Kit (Sigma-Aldrich, St. Louis, MO). DNA concentration and quality were assessed using Nanodrop 1000 spectrophotometer (Thermo Fisher Scientific,Waltham, MA). DNA concentration was converted to number of diploid copies according to the formula: DNA (ng/µl) / weight of mouse diploid genome (3.9 pg). Samples were diluted to 20000 genome copies. The samples were normalized internally according to the number of accepted droplets. Moreover, inter-samples normalization was performed according to the formula (x-min(x))/(max(x)-min(x)) where x represents the accepted droplets. The data were reported as positive droplets / accepted droplets x 100.

Digital droplet PCR protocol and analysis was performed as previously described [50]. Sequences of primers and probe used for the *Kras*$^{Q61R}$ assay are as follows: *Kras*$^{Q61R}$ forward: ATCTGACGTGCTTTGCCTGT, *Kras*$^{Q61R}$ reverse: CCCTCCCCAGTTCTCATGTA, *Kras*$^{Q61R}$ probe: GACACAGCAGGTCAAGAGGAGTACA. The *mT* (*Tomato*) assay is registered as dCNS685684912 (Bio-Rad Laboratories Inc, Hercules, CA) with MIQE context: seq1:195-315:+ CCAGTTCATGTACGGCTCCAAGGCGTACGTGAAGCACCCCGCCGACATCCCCGAT TACAAGAAGCTGTCCTTCCCCGAGGGCTTCAAGTGGGAGCGCGTGATGAACTTCG AGGACGGCGGTCT. Primers and fluorescently labeled probes were combined in a mix containing 18 µM of forward and reverse primers and 5 µM of labeled probes (20x primer / Taqman probe mix).  Reactions were then assembled to contain 12.5 µl of 2x ddPCR mix no-UTP (Bio-Rad Laboratories Inc, Hercules, CA), 1.25 µl of 20x *Kras*$^{Q61R}$ custom Primer / Taqman probe Mix, 1.25 µl of 20x *mT* (*Tomato*) custom Primer / Taqman probe Mix and 10 µl of DNA diluted in nuclease-free water. Digital droplet PCR protocol included a first denaturation step at 95°C for 10 minutes followed by 40 cycles of denaturation at 95°C for 30 seconds and 40 cycles of annealing at 62.5°C for 60 seconds, and was performed in a Biorad Thermal cycler T100. Results were analyzed with a Bio-Rad QX100 droplet reader using the Biorad QuantaSoft software. The amplitude gathering thresholds of positive droplets were set at 3500 for the *mT* probe and at 10000 for the *Kras*$^{Q61R}$ probe, according to the manufacturer's instructions.

**Bronchoalveolar lavage**

Bronchoalveolar lavage (BAL) was performed using three sequential aliquots of 1000 μL sterile ice-cold phosphate-buffered saline (PBS). Fluid was combined and centrifuged at 260 g for 10 min to separate cells from supernatant. The cell pellet was resuspended in 1 ml PBS containing 2% fetal bovine serum, and the total cell count was determined using a grid hemocytometer according to the Neubauer method. Cell differentials were obtained by counting 400 cells on May-Grünwald-Giemsa-stained cytocentrifugal specimens. Total cell numbers in BAL were calculated by multiplying the percentage of each cell type by the total BAL cell number, as described elsewhere [18].

**Bioluminescence imaging**

*R26.Luc;Scgb1a1.Cre* mice, bioluminescent reporters of *Scgb1a1*+ marked cell mass, received one intraperitoneal injection of saline (100 μL saline) or urethane (1g/Kg in 100 μL saline) and were serially imaged before treatment start, and at 150 and 210 days into treatment. Imaging was done on a Xenogen Lumina II (Perkin-Elmer, Waltham, MA) 5-20 min after delivery of 1 mg D-Luciferin sodium in 100 μL of sterile water to the retro-orbital vein, and data were analyzed using Living Image v.4.2 (Perkin-Elmer, Waltham, MA) [18].

**qPCR and microarrays**

Triplicate cultures of $10^6$ LUAD cells, BMDM (obtained by 1-week bone marrow incubation with 100 ng/mL M-CSF), and tracheal AEC (obtained by 1-week incubation of stripped mouse tracheal epithelium in DMEM) were subjected to RNA extraction using Trizol (Thermo Fisher Scientific, Waltham, MA) followed by column purification and DNA removal (Qiagen, Hilden, Germany). Whole lungs were homogenized in Trizol followed by the same procedure. Pooled RNA (5 µg) was quality tested (ABI 2000 bioanalyzer; Agilent Technologies, Sta. Clara, CA), labeled, and hybridized to GeneChip Mouse Gene 2.0 ST arrays (Affymetrix, Sta. Clara, CA). All data were deposited at GEO (http://www.ncbi.nlm.nih.gov/geo/; Accession ID: GSE94981) and were analyzed on the Affymetrix Expression and Transcriptome Analysis Consoles together with previously reported [36-40] murine ATII and human AEC, ATII, AMΦ, non-smokers lung, and LUAD microarray data (Accession IDs: GSE82154, GSE55459, GSE46749, GSE18816, GSE43458). qPCR was performed using first strand synthesis with specific primers (*Scgb1a1*: ATCACTGTGGTCATGCTGTCC and GCTTCAGGGATGCCACATAAC; *Sftpc*: TCGTTGTCGTGGTGATTGTAG and AGGTAGCGATGGTGTCTGCT; *Gusb*: TTACTTTAAGACGCTGATCACC and ACCTCCAAATGCCCATAGTC) and SYBR FAST qPCR Kit (Kapa Biosystems, Wilmington, MA) in a StepOne cycler (Applied Biosystems, Carlsbad, CA). Ct values from triplicate reactions were analyzed with the relative quantification method $2^{-\Delta CT}$relative to *Gusb*.

**Flow cytometry**

BAL cells were suspended in 50 µL PBS with 2% FBS and 0.1% $NaN_3$, were stained with anti-CD45 (#11-0451-85; eBioscience; Santa Clara, CA) and anti-CD11b (#12-0112-82; eBioscience; Santa Clara, CA) primary antibodies for 20 min in the dark at 0.5 µL antibody per million cells, and were analyzed on a CyFlowML cytometer with a sorter module using FloMax Software (Partec, Darmstadt, Germany) or FlowJo software (TreeStar, Ashland, OR), as described previously . Perfused lungs were digested in RPMI-1640 medium containing collagenase XI (0.7 mg/mL; Sigma, St. Louis, MO) and type IV bovine pancreatic DNase (30 µg/mL; Sigma, St. Louis, MO) to obtain single-cell suspensions. After treatment with red blood cell lysis buffer (BioLegend; San Diego, CA), single-cell suspensions were analyzed on a LSR II flow cytometer (BD Bioscience, San Diego, CA), and data were examined with

FlowJo. Dead cells were excluded using 4,6-diamidino-2-phenylindole (DAPI; Sigma, St. Louis, MO).

**Microarray and gene set enrichment analyses (GSEA)**

GSEA was performed with the Broad Institute pre-ranked GSEA module software [http://software.broadinstitute.org/gsea/index.jsp; [51]]. In detail, genes significantly expressed (log2 normalized expression > 8) in murine tracheal airway cells, ATII cells [36], and bone-marrow-derived macrophages were cross-examined against the murine lung and chemical-induced lung adenocarcinoma cell line transcriptomes. In addition, previously reported human airway, ATII, and alveolar macrophage cellular signatures [38-40] were cross-examined against the previously described transcriptomes of human normal lung tissue from never-smokers and of lung adenocarcinomas [37].
