## Supplementary material for "Dual airway and alveolar contributions to adult lung homeostasis and carcinogenesis": Figure S1

### SUPPLEMENTARY FIGURE 1

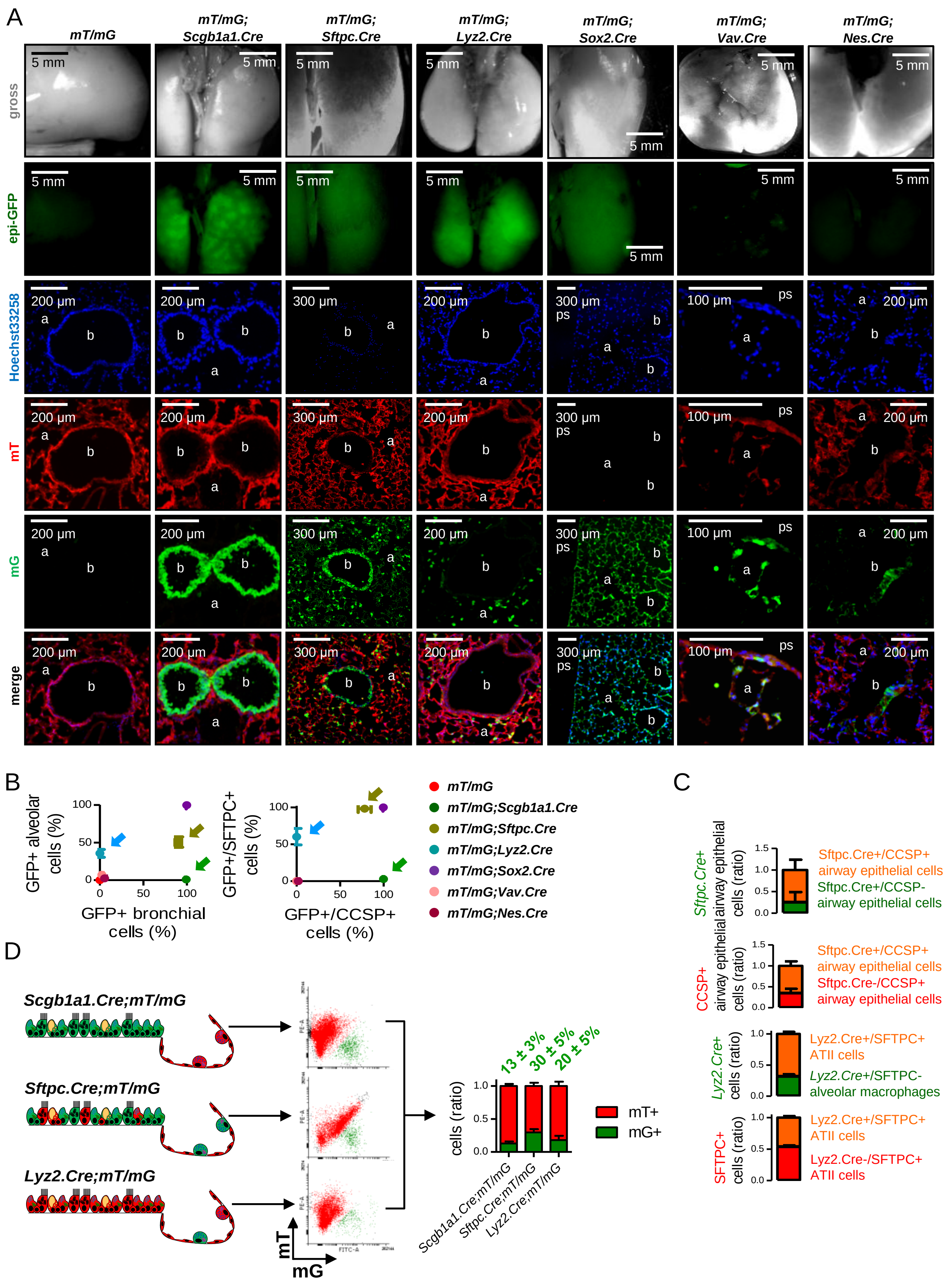

**Supplementary Figure 1.** Mouse models for genetic marking of lung cells. **A,** Lung photographs and green epifluorescence images (top two rows), as well as fluorescent microscopic images of lung sections (Hoechst 33258 stain, endogenous *mT* and *mG* fluorescence, and merged images; bottom four rows) of genetically marked mice employed in these studies at six postnatal weeks (*n* = 5/group). Note absence of *mG*+ cells in *mT/mG* and of *mT*+ cells in *mT/mG;Sox2.Cre* mice, *mG*+ cells in bronchi (b) but not alveoli (a) of *mTmG;Scgb1a1.Cre* mice, in bronchi and alveoli of *mTmG;Sftpc.Cre* mice, in alveoli of *mTmG;Lyz2.Cre* mice, in alveolar capillaries of *mTmG;Vav.Cre* mice, and in neuroepithelial bodies of *mTmG;Nes.Cre* mice. ps, pleural space. **B,** (Left graph), XY plot of *mG*+ airway versus alveolar cells from A (*n* = 5/group). Arrows denote the complete and exclusive *mG*+ marking of airway but not alveolar cells in *mTmG;Scgb1a1.Cre* mice (green); the exclusive marking of some alveolar but not airway cells in *mTmG;Lyz.Cre* mice (blue arrows); and the promiscuous marking of alveolar and airway cells in *mTmG;Sftpc.Cre* mice (olive). (Right graph), data summary from immunostaining of lung sections of lung-marked mice (*n* = 5/group) for CCSP and SFTPC shown in Figure 1A: XY plot of ratios of *mG*+ to CCSP+ airway versus *mG*+ to SFTPC+ alveolar cells. Arrows denote the complete and exclusive *mG*+ marking of CCSP+ club cells, but not of alveolar cells in *mTmG;Scgb1a1.Cre* mice (green); the exclusive marking of a fraction of SFTPC+ ATII cells, but not of airway cells in *mTmG;Lyz.Cre* mice (blue); and the promiscuous marking of *mTmG;Sftpc.Cre* mice (olive). **C,** Quantification of genetic/proteinaceous labeling from immunostains of airways of *mT/mG;Sftpc.Cre* mice (*n* = 5) for Clara cell secretory protein (CCSP, top two graphs) shows that 75 ± 24% of *mG*+ cells are club cells and 66 ± 11% of club cells are *mG*+, and from immunostains of distal alveolar regions of *mT/mG;Lyz2.Cre* mice (*n* = 5) for SFTPC and LYZ2 (bottom two graphs) that 68 ± 3% of *mG*+ cells are ATII cells and 32± 3% alveolar macrophages (AMΦ) and that 47 ± 4% of ATII cells are *mG*+, i.e. from the SFTPC+LYZ2+ lineage. **D,** Schematic representation of genetic marking in *mTmG;Scgb1a1.Cre*, *mTmG;Sftpc.Cre*, and *mTmG;Lyz.Cre* mice (left), flow cytometric gating strategy to quantify *mG*+ and *mT*+ cells (middle), and data summary from *n* = 5, 3, and 6 mice/group (right). Numbers above columns are mean ± SD values of *mG*+ cell percentage/strain. Data are given as mean ± SD. Five non-overlapping fields/sample were examined. *mG*, membranous green fluorescent protein fluorophore; *mT*, membranous tomato fluorophore.
