## Supplementary material for "Dual airway and alveolar contributions to adult lung homeostasis and carcinogenesis": Figure S2

### SUPPLEMENTARY FIGURE 2

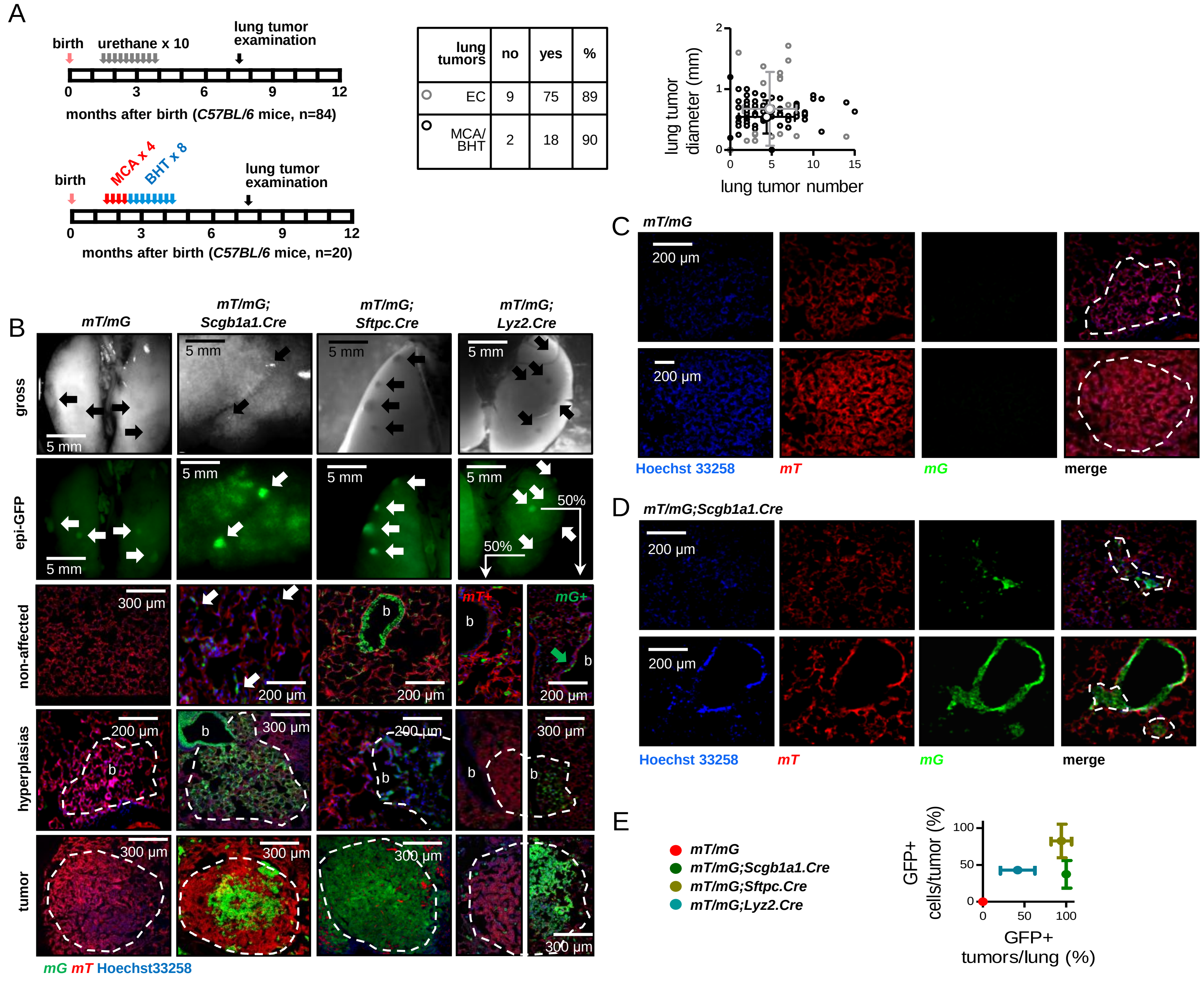

**Supplementary Figure 2.** Changes in pulmonary marked cells during urethane- and MCA/BHT-induced lung adenocarcinoma. **A,** Top left: schematic of multi-hit urethane administration tailored to yield 90% tumor incidence in *C57BL/6* mice: ten weekly intraperitoneal injections of 1 g/Kg urethane (grey arrows) were initiated at six weeks after birth (pink arrow) and lungs were examined six months after the first urethane injection (black arrow). Bottom left: MCA/BHT regimen tailored to yield 90% tumor incidence in *C57BL/6* mice. Four weekly intraperitoneal injections of 15 mg/Kg MCA (red arrows) initiated at six weeks after birth (pink arrow) were followed by eight weekly intraperitoneal injections of 200 mg/Kg BHT (blue arrows) and lung examination at six months after first MCA (black arrow). Lung tumor incidence, number and mean diameter are depicted in table and plot, respectively. **B,** Photographs and green epifluorescence images of tumor-bearing lungs (top two rows), as well as merged fluorescent microscopic images of lung sections (of Hoechst 33258 stain and endogenous *mT* and *mG* fluorescence) taken from non-affected, hyperplastic/proliferative and frank neoplastic regions (bottom three rows) from experiment from Figure 1C (*n* = 30, 22, 18, and 20/group, respectively). Top two rows: arrows indicate lung tumors. Note the absence of *mG*+ fluorescence in *mT/mG* tumors, the *mG*+ fluorescence of *mT/mG;Scgb1a1.Cre* and *mT/mG;Sftpc.Cre* tumors, and the split *mG*+ and *mG*- tumors of *mTmG;Lyz2.Cre* mice. Bottom three rows: white arrows indicate *mG*+ cells in apparently non-affected alveolar areas of *mT/mG;Scgb1a1.Cre* mice; green arrow indicates rare *mG*+ cell in non-affected airway of *mT/mG;Lyz2.Cre* mouse. Note absence of *mG*+ fluorescence in *mT/mG* hyperplasias and tumors (dashed outlines), *mG*+ fluorescence of *mT/mG;Scgb1a1.Cre* and *mT/mG;Sftpc.Cre* lesions, and split *mG*+ and *mG*- lesions of *mTmG;Lyz2.Cre* mice. **C** and **D,** Single-channel (endogenous *mT* and *mG* fluorescence and Hoechst 33258 nuclear stain) and merged images of lung neoplasias (dashed outlines) of *mT/mG* and *mT/mG;Scgb1a1.Cre* mice treated with MCA/BHT as in A (*n* = 8/group). Note absence of *mG*+ fluorescence in *mT/mG* tumors and the *mG*+ tumors of *mT/mG;Scgb1a1.Cre* mice. **E,** XY plot of percentage of *mG*+ tumors/lung versus *mG*+ tumor cells/tumor averaged per lung from B (*n* = 30, 22, 18, and 20/group, respectively). Five non-overlapping fields/lung were examined. Data are given as mean ± SD. *mG*, membranous green fluorescent protein fluorophore; *mT*, membranous tomato fluorophore.
