## Supplementary material for "Dual airway and alveolar contributions to adult lung homeostasis and carcinogenesis": Figure S3

SUPPLEMENTARY FIGURE 3

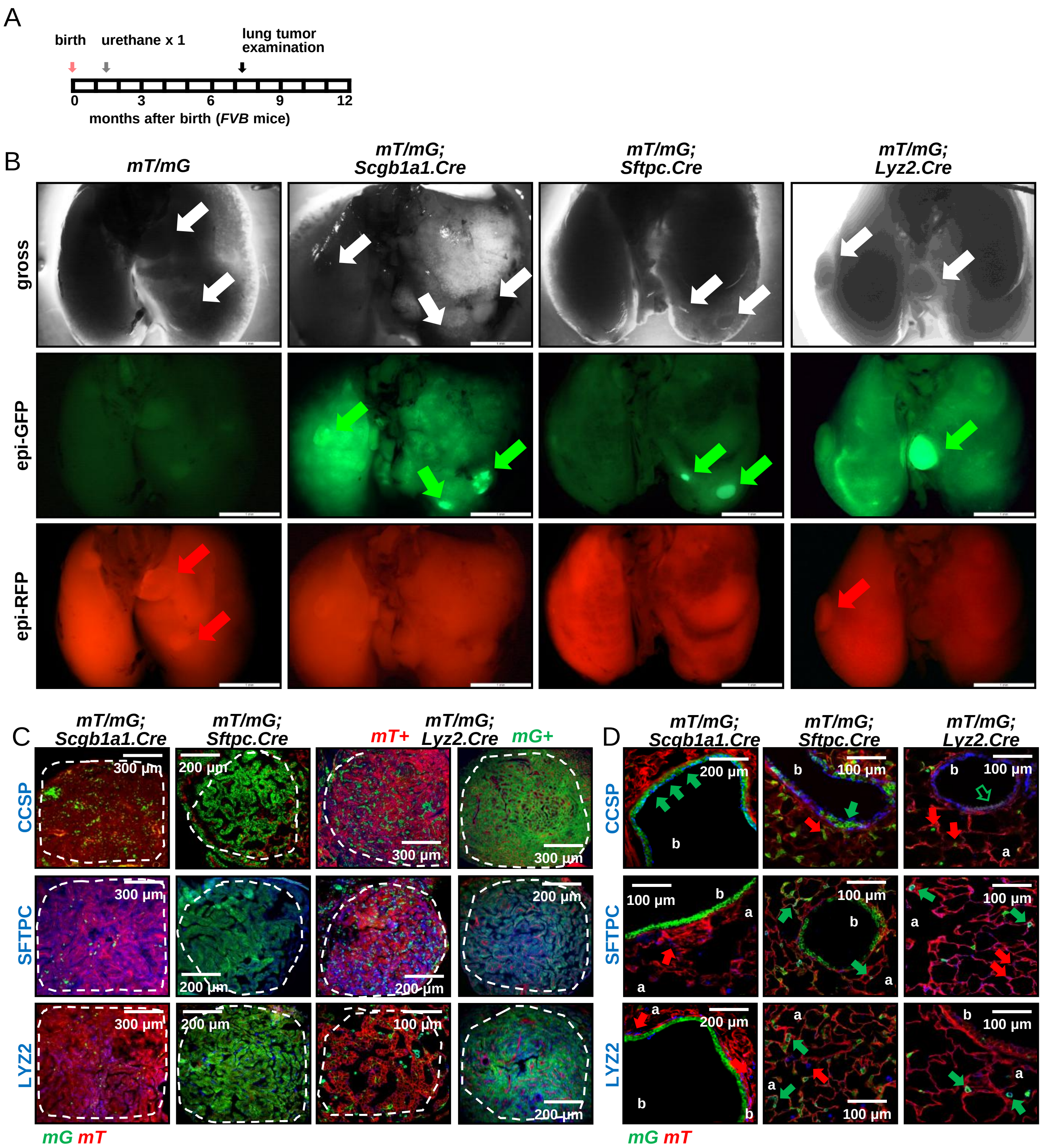

**Supplementary Figure 3.** Lung adenocarcinoma molecular signatures of *FVB* mice. **A,** Schematic of urethane administration in *FVB* mice: one intraperitoneal injection of 1 g/Kg urethane (grey arrow) was administered at six weeks after birth (pink arrow) and lungs were examined six months later (black arrow). **B,** Representative photographs and epifluorescence images of tumor-bearing *mT/mG*, *mT/mG;Scgb1a1.Cre*, *mT/mG;Sftpc.Cre*, and *mTmG;Lyz2.Cre FVB* lungs (*n*≥ 8/group). Arrows indicate lung tumors. Note the absence of *mG*+ fluorescence in *mT/mG* tumors, the *mG*+ fluorescence of *mT/mG;Scgb1a1.Cre* and *mT/mG;Sftpc.Cre* tumors, and the split *mG*+ and *mG*- tumors of *mTmG;Lyz2.Cre* mice. **C,** Lineage marker protein-stained LUAD (dashed outlines) from genetically-marked mice from D (*FVB, n*≥ 10/group). Note *mG*+CCSP-SFTPC+LYZ2± LUAD cells of *mTmG;Scgb1a1.Cre* mice. **D,** Lineage marker-stained lung sections of 6-week-old lung-marked mice (*FVB*, *n* = 5/group). Note *mG*+CCSP+ club and *mG*+TUBA1A+ ciliated cells in bronchi (b) of *mT/mG;Scgb1a1.Cre* mice, *mG*+SFTPC+LYZ2± alveolar type II cells and *mG*+SFTPC-LYZ2+ alveolar macrophages in the alveoli (a) of *mT/mG;Lyz2.Cre* mice, and various *mG*+ cells in bronchi and alveoli of *mT/mG;Sftpc.Cre* mice. Arrows indicate lineage marker protein-expressing *mG*+ (green) and *mT*+ (red) cells. Five non-overlapping fields/sample were examined. *mG*, membranous green fluorescent protein fluorophore; *mT*, membranous tomato fluorophore; CCSP, Clara cell secretory protein; SFTPC, surfactant protein C; LYZ2, lysozyme 2.
