## Supplementary material for "Dual airway and alveolar contributions to adult lung homeostasis and carcinogenesis": Figure S4

### SUPPLEMENTARY FIGURE 4

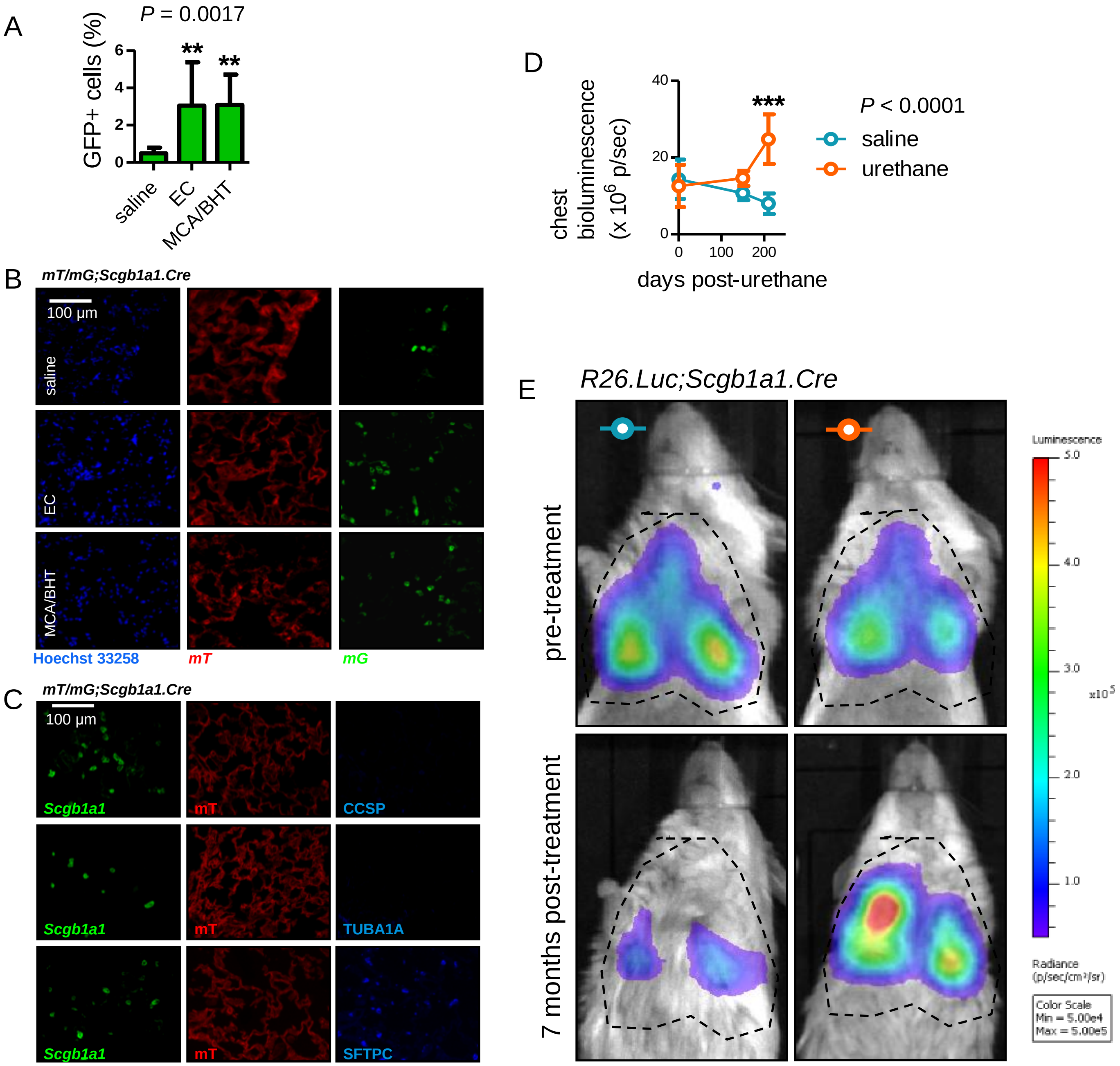

**Supplementary Figure 4.** Expansion of *Scgb1a1*+ marked cells during urethane- and MCA/BHT-triggered field cancerization of the lungs. **A** and **B,** Data summary of alveolar *mG*+ cells (A) and single-channel (endogenous *mT* and *mG* fluorescence and Hoechst 33258 nuclear stain) images (B) corresponding to Figure 3A, showing non-neoplastic alveolar regions of saline-, EC-, and MCA/BHT-treated *mT/mG;Scgb1a1.Cre* mice at six months into treatment (*n* = 8 mice/group). Note the few *mG*+ cells of saline-treated mice and increased numbers in carcinogen-treated mice. **C,** Single-channel images corresponding to Figure 3B. Shown are non-neoplastic distal lung regions of EC-treated *mT/mG;Scgb1a1.Cre* mice at six months into treatment (n = 22), stained for the lung cell markers Clara cell secretory protein (CCSP), acetylated α-tubulin (TUBA1A), and surfactant protein C (SFTPC). Note *mG*+CCSP-TUBA1A1-SFTPC+ cells in the alveoli. **D** and **E,** Data summary (D) and representative bioluminescence images (merged with photographic images; E) of *R26.Luc;Scgb1a1.Cre FVB* mice before and seven months after saline (one intraperitoneal injection of 100 μL; *n* = 6) or EC (one intraperitoneal injection of 1 g/Kg in 100 μL saline; *n* = 5) treatment. Note that in this model light is emitted exclusively by *Scgb1a1*+ marked cells. Note also that this signal was exclusively detected over the lungs and over the female pelvis during the productive phase of the menstrual cycle, emitted by endometrial cells that also express *Scgb1a1* (data not shown). Note the ~30% reduction in the signal of saline-treated mice over the seven months of the experiment that is likely attributable to reduced light penetrance through the thoracic wall of the growing mouse and not to reduction of *Scgb1a1*+ marked cells. Note the ~30% increase in the signal of EC-treated mice over the same time period that integrally shows expansion of *Scgb1a1*+ cells in response to EC. Measurements were from five non-overlapping fields/lung. Data are given as mean ± SD. **, and ***: $P< 0.01$, and $P< 0.001$ for comparison of the indicated columns with saline by one-way ANOVA with Bonferroni post-tests (A) and of urethane and saline treatments at the 7-month-time-point by two-way repeated measures ANOVA with Bonferroni post-tests (D). *mG*, membranous green fluorescent protein fluorophore; *mT*, membranous tomato fluorophore.
