## Supplementary material for "Dual airway and alveolar contributions to adult lung homeostasis and carcinogenesis": Figure S5

### SUPPLEMENTARY FIGURE 5

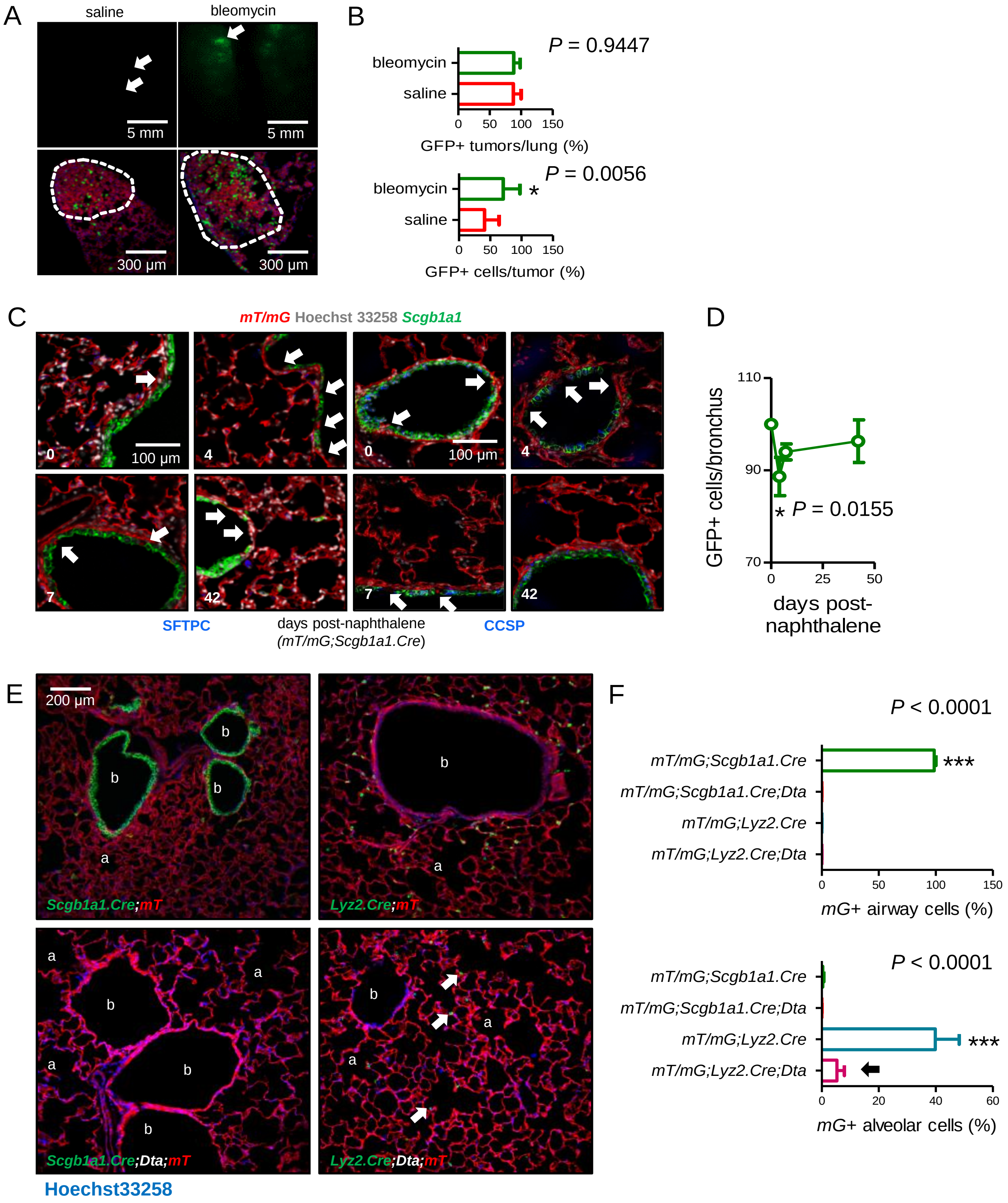

**Supplementary Figure 5.** *Scgb1a1*+ marked cells in lung injury and triple transgenic models of pulmonary lineage ablation. **A** and **B,** Representative epifluorescence images of tumor-bearing lungs (A; top), as well as merged fluorescent microscopy images of lung tumors (merges of endogenous *mT* and *mG* stained with Hoechst 33258; A, bottom) and data summary of *mG*+ tumors and tumor cells (B) of six-week-old *mT/mG;Scgb1a1.Cre* mice that received saline (*n* = 12) or 0.08 units bleomycin (*n* = 13) intratracheally, were allowed to recover for one month, and subsequently received ten weekly intraperitoneal injections of 1 g/Kg urethane to be sacrificed six months after first urethane injection. Arrows and dashed outlines in (A) indicate tumors. Note the enrichment of lung adenocarcinomas in *mG*+ cells in response to bleomycin, which depletes resident alveolar type II (ATII) cells, as shown in Figure 4B. **C** and **D,** Representative fluorescent microscopic images [C; merges of Hoechst 33258-stained endogenous *mT* and *mG* immunostained for surfactant protein C (SFTPC, left) or Clara cell secretory protein (CCSP, right)] and data summary of percentage of *mG*+ airway cells (D; *n* = 5 mice/group) of *mT/mG;Scgb1a1.Cre* mice after intraperitoneal injection of 250 mg/Kg naphthalene given at six weeks of age. Arrows denote naphthalene-induced airway epithelial gaps that are restored by *mG*+CCSP+SFTPC- cells. **E** and **F,** Representative lung sections of 12-week-old *mT/mG;Scgb1a1.Cre*, *mT/mG;Lyz2.Cre*, *mT/mG;Scgb1a1.Cre;Dta*, and *mT/mG;Lyz2.Cre;Dta* mice (E; *n* = 6/group) and data summary of airway (top) and alveolar (bottom) *mG*+ cells (F; *n* = 6/group). Note increased bronchial (b) and alveolar (a) size, complete airway epithelial denudement, and prominent distortion of bronchial and alveolar structure of *mT/mG;Scgb1a1.Cre;Dta* mice compared with other strains, mimicking human emphysema. Note also *mG*+ alveolar macrophages (AMΦ) in *mT/mG;Lyz2.Cre;Dta* mice (arrows). Measurements were from at least five non-overlapping tumor, airway, or alveolar fields/lung. Data are given as mean ± SD. ns, *, and ***: $P > 0.05$, $P < 0.05$, and $P < 0.001$, respectively, for the indicated comparisons by unpaired Student's t-test (B), for comparison with time-point zero by one-way ANOVA with Bonferroni post-tests (D), or for comparison of the indicated columns to all other groups by one-way ANOVA with Bonferroni post-tests (F). *mG*, membranous green fluorescent protein fluorophore; *mT*, membranous tomato fluorophore.
