## Supplementary material for "Dual airway and alveolar contributions to adult lung homeostasis and carcinogenesis": Figure S6

### SUPPLEMENTARY FIGURE 6

A

| | | Signature genes | |
|---|---|---|---|
| | | Lung | LADC |
| Signature genes | AEC | 23 | 25 |
| | ATII | 24 | 10 |
| | BMDM | 14 | 9 |

AEC vs ATII: * ; ATII vs BMDM: ns ; AEC vs BMDM: ns

$P$ = 0.1168

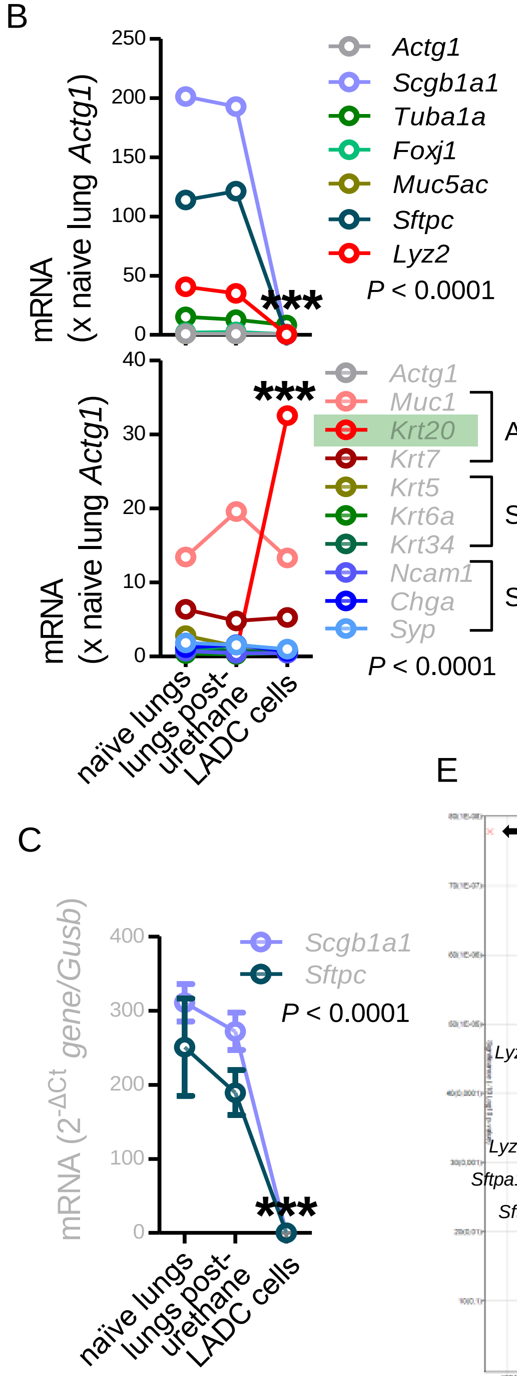


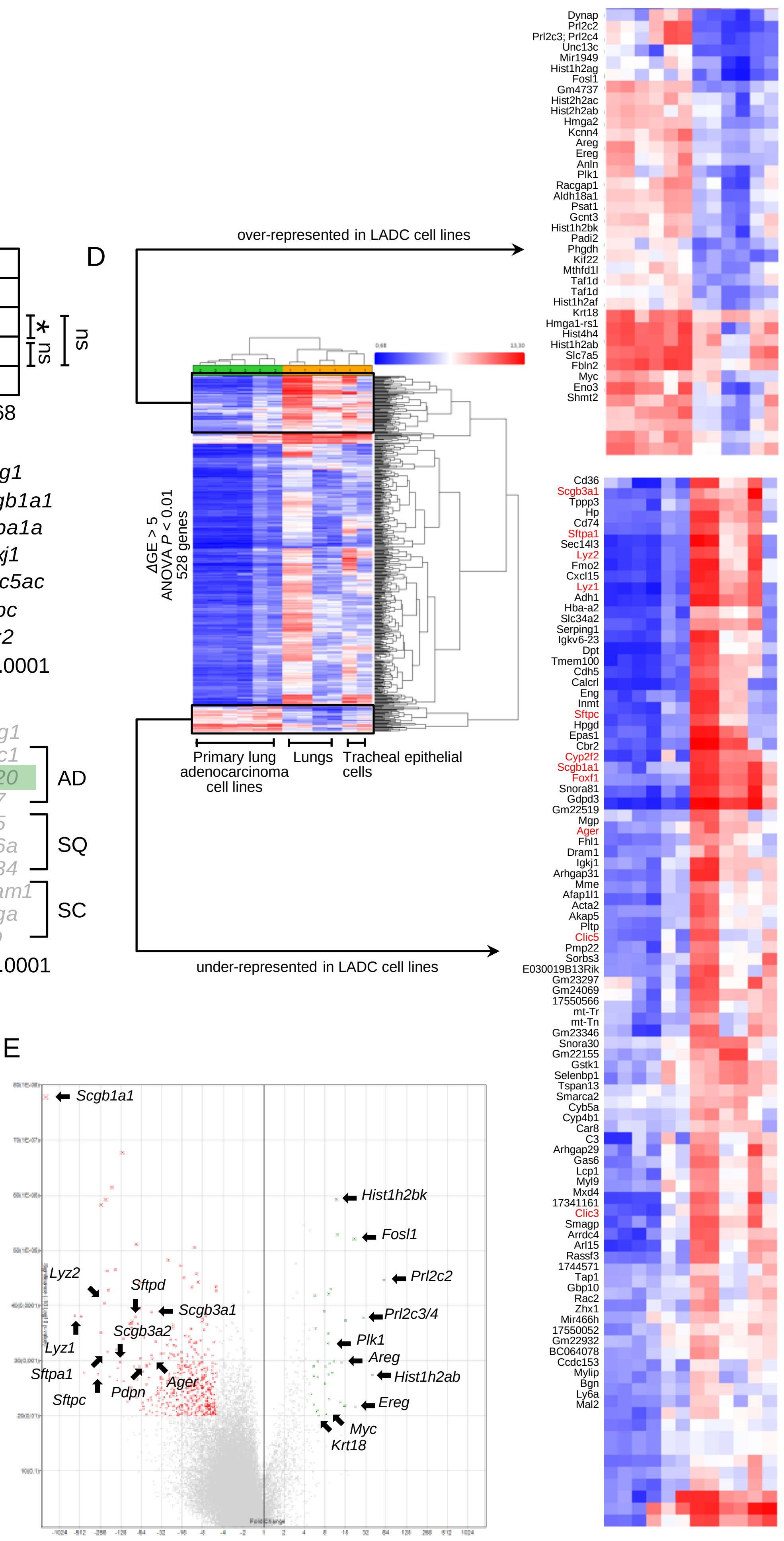

**Supplementary Figure 6.** Transcriptomic characterization of urethane-induced lung adenocarcinoma (LUAD) cell lines. **A-C,** RNA of mouse urethane-induced LUAD cell lines, lungs obtained pre- and one week post-urethane treatment (*n* = 2 each), and airway epithelial cells (AEC), alveolar type II cells [ATII; data from [36]], and bone marrow-derived macrophages (BMDM) was examined by Affymetrix Mouse Gene ST2.0 microarrays (*n* = 4/group). **A,** Number of genes out of the 30 top-represented transcripts of AEC, ATII, and BMDM within the top-2000-expressed genes of lungs and LUAD cells. **B** and **C,** Mean expression levels of selected transcripts, including lineage markers and markers of histologic subtype in LUAD compared with lungs pre- and one week post-urethane treatment (B, microarray, *n* = 2/group; C, qPCR, *n* = 3/group). AD, adenocarcinoma; SQ, squamous cell carcinoma; SC, small cell carcinoma. **D** and **E,** Differentially expressed genes between six different lung adenocarcinoma cell lines cultured from urethane-induced lung tumors and six benign respiratory mouse samples, including lungs of saline- and urethane-treated mice obtained at one week post-treatment, as well as primary mouse tracheal epithelial cells. **D,** Whole heat map showing the accurate hierarchical clustering of the samples according to differentially expressed genes, as well as the top over- and under-represented genes. Note the universal loss of expression of lineage markers by lung adenocarcinoma cells (genes in red font). **E,** Volcano plot showing selected top over- and under-represented genes (arrows). ANOVA, analysis of variance; FDR, false discovery rate. Data are given as mean ± SD and two-way ANOVA (graphs) or $\chi^2$ test (table) *P* values. ns, *, and ***: *P*> 0.05, *P*< 0.05, and *P*< 0.001 for the comparisons indicated (A) or for comparison of all genes (B, D) and of *Krt20* (C) with naïve lungs by Bonferroni post-tests (graphs) or Fischer's exact test (table).
