## Supplementary material for "Dual airway and alveolar contributions to adult lung homeostasis and carcinogenesis": Figure S7

SUPPLEMENTARY FIGURE 7

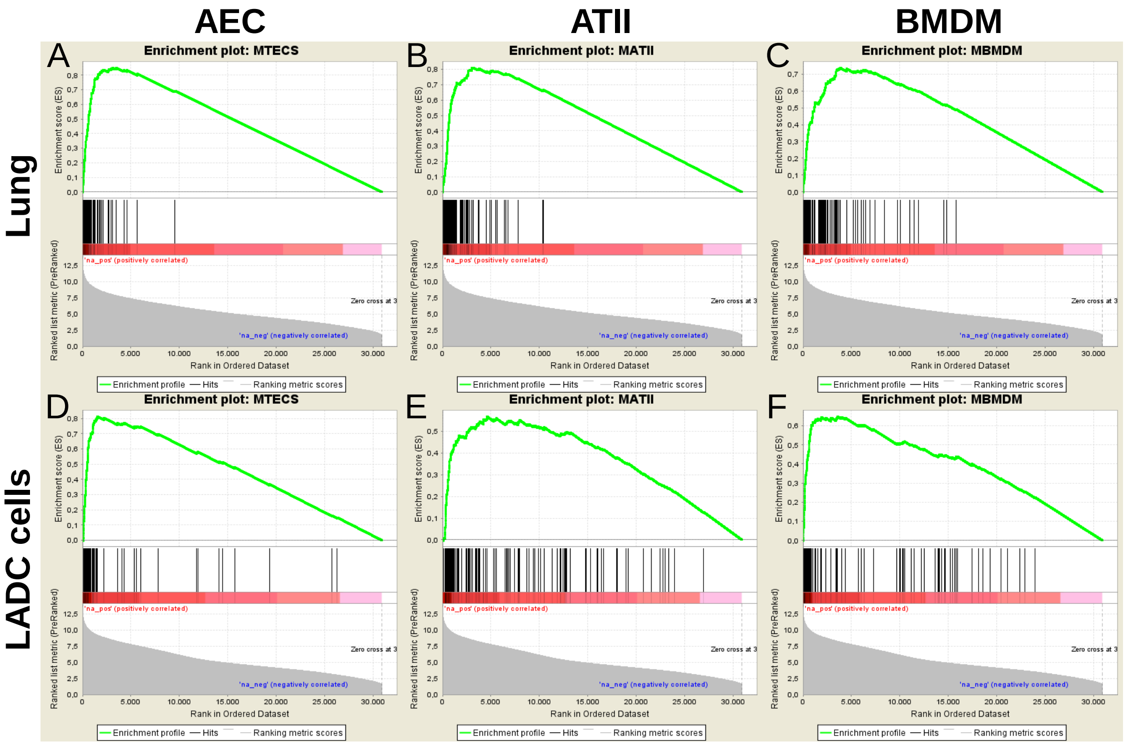


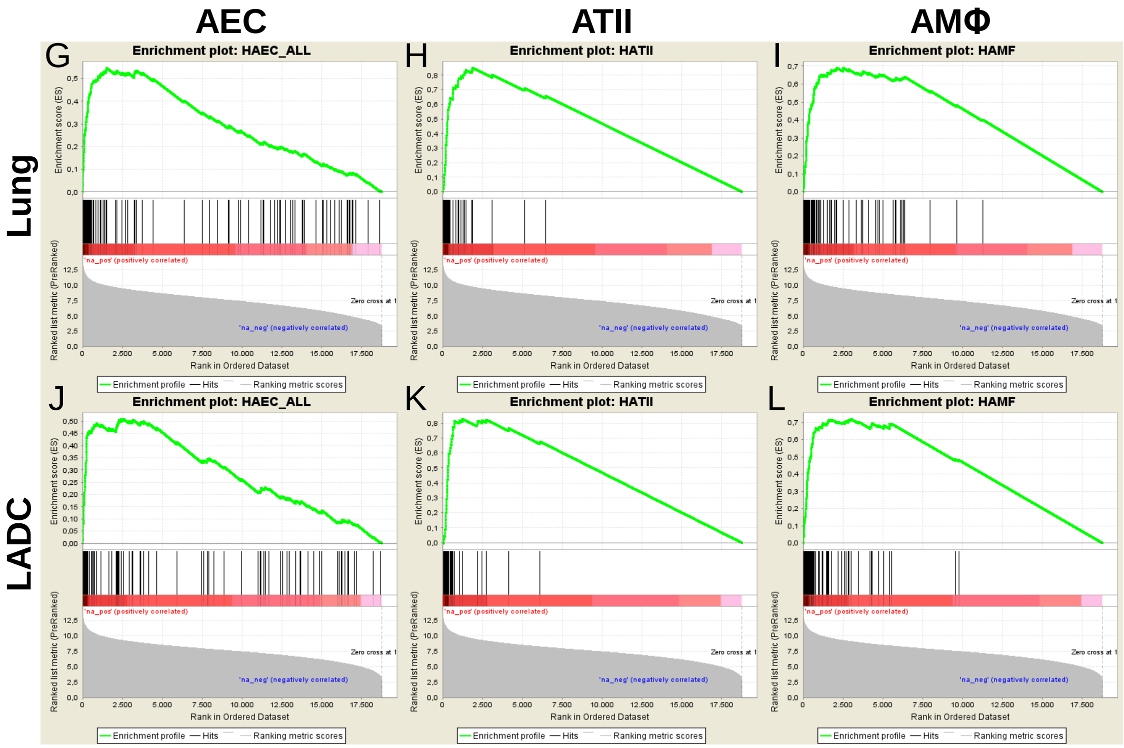

**Supplementary Figure 7.** Transcriptomic characterization of urethane-induced lung adenocarcinoma (LUAD) cell lines. **A-F,** RNA of mouse urethane-induced LUAD cell lines, lungs obtained pre- and one week post-urethane treatment (*n* = 2 each), airway epithelial cells (AEC), alveolar type II cells [ATII; data from [36]], and bone marrow-derived macrophages (BMDM) was examined by Affymetrix Mouse Gene ST2.0 microarrays (*n* = 4/group). Gene set enrichment analysis, including enrichment scores and nominal probability values, of AEC, ATII, and BMDM signatures in mouse LUAD cell transcriptome shows significant enrichment of the AEC (but not the ATII and BMDM/AMΦ) signature in LUAD compared with lung. **G-L,** RNA of human LUAD (*n* = 40), never-smoker lung tissue (*n* = 30), primary AEC (*n* = 5), primary ATII (*n* = 4), and alveolar macrophages (AMΦ; *n* = 9) analyzed by Affymetrix Human Gene ST1.0 microarrays was cross-examined [data from [37-40]]. Gene set enrichment analysis, including enrichment scores and nominal probability values, of AEC, ATII, and AMΦ signatures in LUAD transcriptome shows significant enrichment of the AEC (but not the ATII and BMDM/AMΦ) signature compared with lung. Nominal $P< 0.0001$ for all comparisons, family-wise error rates FWER < 0.01.
